# Huntingtin preserves mitochondrial genome integrity in neurons, which is impaired in Huntington’s disease

**DOI:** 10.1101/2025.07.24.666629

**Authors:** Subrata Pradhan, Sagar Gaikwad, Chi-Lin Tsai, Charlene Smith, Nan Zhang, Keegan Bush, Anirban Chakraborty, Subo Yuan, Sanjeev Choudhary, C. Dirk Keene, Lisa M. Ellerby, Tapas K. Hazra, Albert R. La Spada, Yogesh P. Wairkar, Tatsuo Ashizawa, John A. Tainer, Tej K. Pandita, Leslie M. Thompson, Partha S. Sarkar

## Abstract

Huntingtin (HTT) function is enigmatic, as the native protein plays critical roles in neuronal health, while mutant HTT (mHTT), carrying an expanded polyglutamine stretch, triggers neurotoxicity and contributes to the pathogenesis of Huntington’s disease (HD). We recently found that HTT is part of a nuclear transcription-coupled DNA repair (TCR) complex with DNA repair enzymes including polynucleotide-kinase-3’-phosphatase (PNKP). This complex resolves DNA lesions during transcription to maintain genome integrity, while in HD, mHTT impairs the activity of this complex, resulting in accumulation of DNA lesions. Using molecular, cellular biology and computational methods, we find that HTT has a role in assembling a functional DNA repair complex in mitochondria. Together with mitochondrial RNA polymerase and transcription factors, HTT resolves mitochondrial DNA lesions to preserve mitochondrial genome integrity and function. Pathogenic mHTT impairs this activity, resulting in persistent DNA lesions and reduced mitochondrial function in HD. Importantly, restoring activity of this complex in a *Drosophila* HD model through ectopic HTT or PNKP expression significantly improves mitochondrial genome integrity and ameliorates motor deficits.

**HIGHLIGHTS:**

- HTT organizes a functional, multifactorial mitochondrial DNA repair complex
- Mutant HTT impairs the mitochondrial DNA repair complex causing DNA damage accumulation
- HTT-associated repair complex resolves mitochondrial DNA lesions and DNA integrity
- Restoring repair activity in HD flies rescues mitochondrial DNA integrity and motor defects

## INTRODUCTION

Huntingtin (HTT) is a 3114 amino acid protein that functions have a wide range of roles in neuronal health. Wild-type HTT (wtHTT) protects cells against pro-death stimuli, while HTT depletion increases cellular vulnerability[1–5]. Consistent with these observations, transgenic wtHTT expression in mouse brain confers neuroprotection against excitotoxic and ischemic injury[6] whereas brain-specific HTT inactivation triggers neurodegeneration and motor deficits[7]. Recent studies indicate HTT’s role in the protection of mitochondrial function[8]. However, the mechanism by which wtHTT confers neuroprotection is unclear.

Huntington’s disease (HD) is a dominantly inherited fatal neurodegenerative disease caused by CAG repeat expansion in the *HTT* gene, encoding an extended polyglutamine (polyQ) sequence in mutant HTT (mHTT)[9]. Mutant HTT (mHTT) gains a toxic function, and disrupts mitochondrial function, thus contributing to HD pathology. For instance, mHTT associates with the outer mitochondrial membrane and impairs mitochondrial protein import[10–12] and disrupts mitochondrial quality control through dynamin-related protein-1 (DRP1)[13]. Additionally, mitochondrial DNA (mtDNA) damage is observed in HD and implicated in perturbing energy homeostasis thus causing mitochondrial dysfunction[14–18]. These findings indicate that mitochondrial dysfunction significantly contributes to HD progression.

Consistent with the postulated role of wtHTT in facilitating the assembly of multiprotein complexes[19, 20], we recently demonstrated that wtHTT assembles a macromolecular transcription-coupled DNA repair (TCR) complex with RNA polymerase II subunit A (POLR2A), ataxin-3, and essential DNA repair enzymes including DNA ligase 3 (Lig3), and polynucleotide-kinase-3’-phosphatase (PNKP) in the nucleus[21]. This HTT-assembled multifactorial complex helps in maintaining the genome integrity and neuronal function by resolving the DNA damage that occur during transcription[21]. However, mHTT impaired the nuclear TCR complex’s DNA repair activity, resulting in persistent DNA damage accumulations in HD and various models of HD[21]. While this nuclear repair complex is well-established, it remains unknown whether HTT assembles a similar mitochondrial DNA repair complex to address mtDNA damage.

The mitochondrial genome is constantly exposed to reactive oxygen species (ROS) generated by the nearby electron transport chain (ETC) complexes. Importantly, mtDNA lacks protective nucleosomes, making it highly vulnerable to ROS-induced damage[22]. For these reasons, a range of DNA repair mechanisms have evolved to resolve mtDNA damage and to maintain mtDNA integrity[23]. Base excision repair (BER) is the primary mechanism for repairing the oxidized mtDNA bases, but there is some evidence to support nucleotide excision repair (NER) within the mitochondria[23]. Knockdown of wtHTT in murine embryonic stem cells disrupts the structural and functional integrity of mitochondria[8], emphasizing its importance in maintaining mitochondrial integrity. Finally, previous studies have implicated a role for mHTT in mitochondrial degeneration and dysfunction[14, 15, 18, 24]. While these data suggest that HTT may play an important role in maintaining mitochondrial function, precisely how this is achieved and how mHTT disrupts mitochondrial function is not yet understood.

Here, we characterize a functional mtDNA repair complex analogous to our previously described nuclear DNA repair complex[21]. This novel multiprotein complex is composed of HTT, mitochondrial RNA polymerase (POLRMT), mitochondrial DNA polymerase gamma, Cockayne syndrome proteins, PNKP and other mitochondria-specific transcription factors. Importantly, we show that increasing HTT levels in a neuronal cell and *Drosophila* model of HD dramatically improves mtDNA integrity against stress-induced mtDNA damage. Consistent with its role in mtDNA repair, reduction of wtHTT levels or mHTT expression adversely affects the ability of mtDNA to repair itself and decreases repair function. By characterizing this HTT DNA repair complex in mitochondria, we postulate a potential mechanism of how depletion of endogenous HTT disrupts the structural and functional integrity of mitochondria and why and how wtHTT might provide neuroprotection against stress-mediated injury. Together our findings explain how mHTT may cause early mitochondrial dysfunction and neurotoxicity in HD.

## RESULTS

### HTT associates with mitochondrial RNA polymerase, transcription factors, and key DNA repair proteins within mitochondria

Given that our previous studies showed the importance of the wtHTT in stimulating transcription coupled non-homologous end joining DNA damage repair in the nucleus[21] and that HTT appears to confer a protective role to mitochondria[25, 26], we hypothesized that native HTT may protect mitochondria by facilitating mtDNA damage repair. To investigate this possibility, we first assessed the presence and expression levels of HTT, PNKP, and ATXN-3, which are integral components of the nuclear transcription-coupled DNA repair complex that we recently described[21]. We conducted cell fractionation experiments and verified fraction purity using specific markers: GAPDH (cytosolic) fraction: POLR2A (nuclear); POLRMT (mitochondrial). After purifying the mitochondrial (ME), nuclear (NE), and cytosolic (CE) protein fractions, western blotting analysis confirmed HTT and PNKP presence across subcellular fractions (Figure 1A). Consistent with our hypothesis, we observed the presence of both proteins in mitochondrial and nuclear fractions (Figure 1A, arrows), suggesting that HTT and PNKP are associated with mitochondria. To further confirm HTT’s mitochondrial presence, we analyzed the SH-SY5Y cells by immunostaining with an anti-HTT antibody, followed by electron microscopy. The immuno electron microscopic analysis confirmed the presence of native HTT within mitochondria (Figure 1B, arrows).

**Figure 1:**
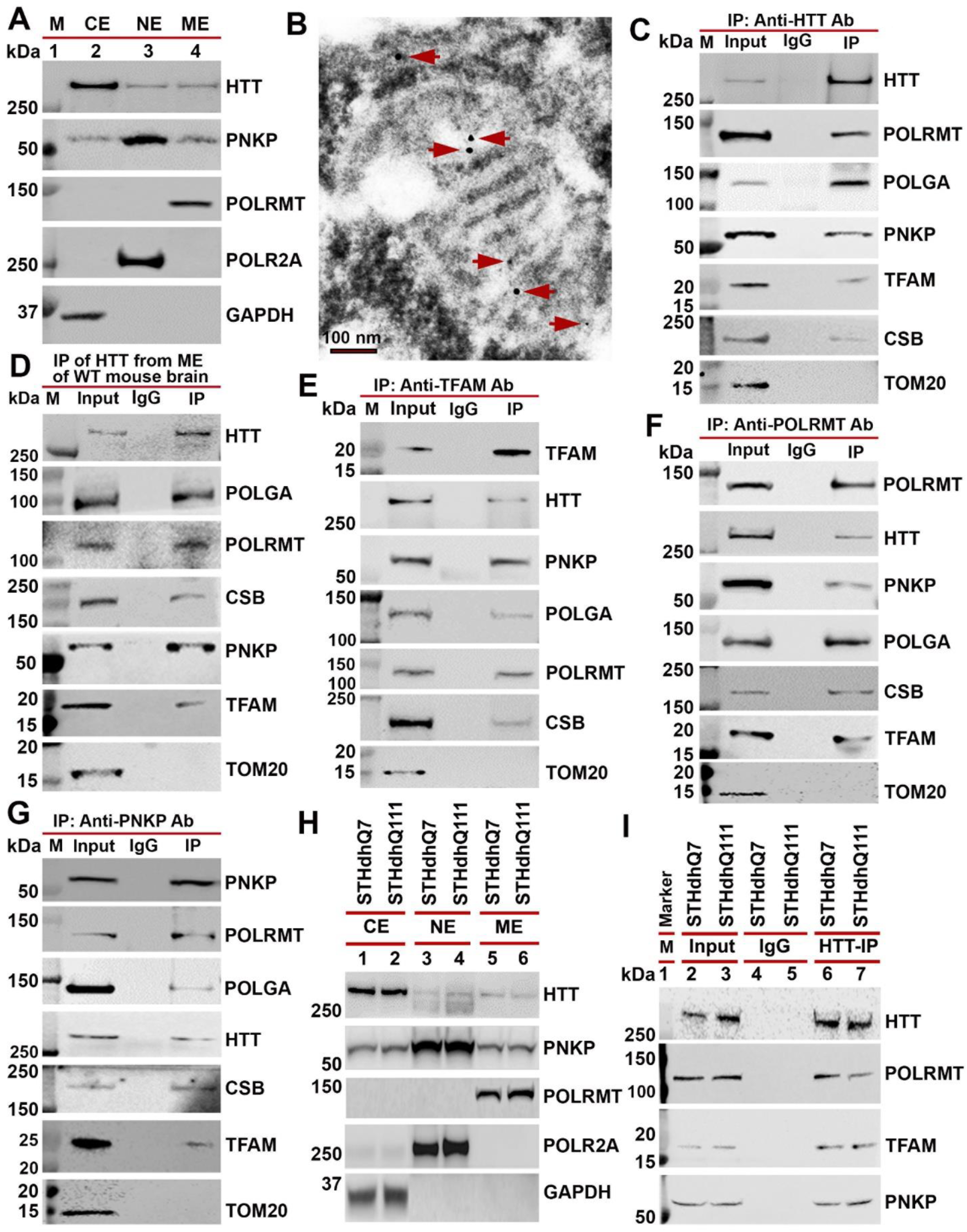
HTT associates with mitochondrial transcription and DNA repair complex. **(A).** Cytosolic, nuclear, and mitochondrial protein extracts (CE, NE, and ME, respectively) are isolated from human neuroblastoma SH-SY5Y cells and these subcellular fractions analyzed by western blotting (WBs) to detect HTT, and PNKP in the fractions. GAPDH (Glyceraldehyde-3-phosphate dehydrogenase), POLR2A (RNA polymerase II large subunit A) and POLRMT (Mitochondrial RNA polymerase) used as cytosolic, nuclear, and mitochondrial protein markers, respectively. Lane 1: M: Protein molecular weight markers in kDa. **(B).** HTT is present within the mitochondria. SH-SY5Y cells were analyzed by immunostaining with anti-HTT primary antibody and gold-coated mouse secondary antibody followed by transmission electron microscopic analysis. The presence of HTT within the mitochondria is shown by arrows. **(C).** Mitochondrial protein extract (MEs) isolated from SH-SY5Y cells, and the HTT IP’d with anti-HTT mouse monoclonal Ab (MAB2170; Millipore-Sigma) from the MEs and HTT immunocomplexes (ICs) subjected to WBs to detect the DNA repair associated proteins. **(D).** MEs isolated from 3-month-old wildtype C57BL/6 mouse cortex tissue (n=3), and the endogenous HTT IP’d with an anti-HTT mouse monoclonal antibody (MAB2170; Millipore-Sigma). The HTT ICs analyzed by WBs to check the presence of DNA repair complex proteins *in vivo*. TOM 20 was used as negative control for C, D, E and F. **(E).** MEs isolated from wildtype C57BL/6 mouse cortex, and the TFAM IP’d with an anti-TFAM rabbit monoclonal Ab (ab 176558; Abcam) from MEs and TFAM ICs analyzed by WBs to detect the mtDNA repair complex-associated proteins. **(F).** MEs purified from wildtype C57BL/6 mouse cortex, and mitochondrial RNA polymerase (POLRMT) IP’d with an anti-POLRMT antibody (GTX105137; Genetex). The POLRMT ICs analyzed by WBs to check the presence of DNA repair complex proteins (HTT, PNKP, POLGA, POLRMT, CSB, and TFAM). **(G).** MEs isolated from the wildtype C57BL/6 mouse cortex, and endogenous PNKP IP’d with anti-PNKP rabbit Ab (NBP1-87257; Novus) from MEs and the PNKP ICs subjected to WBs to detect the same DNA repair complex-associated protein components. **(H).** Cytosolic, nuclear, and mitochondrial protein extracts (CE, NE and ME extracts respectively) were isolated from wildtype STHdhQ7 (lanes 1, 3 and 5) and mutant STHdhQ111 (lanes 2, 4 and 6) mouse striatal cells, and the protein fractions analyzed by WBs to detect HTT, and PNKP in the subcellular fractions. GAPDH (Glyceraldehyde-3-phosphate dehydrogenase), POLR2A (RNA polymerase II subunit A) and POLRMT (Mitochondrial RNA polymerase) used as cytosolic, nuclear, and mitochondrial protein markers, respectively. **(I).** MEs isolated from STHdhQ7 and STHdhQ111 striatal cells, and endogenous HTT immunoprecipitated with an anti-HTT Ab (MAB2170; Millipore-Sigma), and the HTT ICs analyzed by WBs to detect HTT, POLRMT, TFAM and PNKP in the HTT IC. Lane 1: Protein molecular weight marker. Lanes 2 and 3: Input from STHdhQ7 and STHdhQ111 cells respectively, Lanes 4 and 5: IgG IP from STHdhQ7 and STHdhQ111 cells respectively, and lanes 6 and 7: HTT IP from STHdhQ7 and STHdhQ111 cells respectively.

Next, we investigated whether mtDNA repair is also carried out concurrently with transcription given our findings that the nuclear DNA repair mediated by HTT and PNKP is dependent on transcription. Specifically, we tested whether wtHTT associates with the mitochondrial RNA polymerase (POLRMT), mitochondria-specific DNA polymerase (DNA polymerase γ) and mitochondrial transcription factors. We purified mitochondria from SH-SY5Y cells and removed non-specific proteins using protease digestion to prevent contamination that might confound our analyses[27]. Western blotting analysis using nuclear (POLR2A) and cytosolic (GAPDH) protein markers showed no contaminating bands, confirming that mitochondrial fractions were pure (Figure 1A). We reasoned that if wtHTT plays a role in mtDNA repair during transcription, it would likely interact with mtDNA and with mtDNA repair proteins such as PNKP, and the essential components of the known mitochondrial transcription complex, namely, POLRMT, POLGA (catalytic subunit of mitochondria-specific DNA polymerase γ), and mitochondrial transcription factor TFAM (mtTFA or TFAM).

To test these interactions, we used the purified mitochondrial protein extract (MEs) from SH-SY5Y cells and performed immunoprecipitation (IP) studies using antibodies against HTT, POLRMT, PNKP and TFAM. We first immunoprecipitated (IP’d) wtHTT from the MEs isolated from SH-SY5Y cells using an anti-HTT antibody under stringent IP conditions to ensure that we were not immunoprecipitating any non-specific proteins. WB analysis revealed the presence of endogenous POLRMT, PNKP, POLGA, PNKP, and TFAM in the wtHTT immunocomplexes (ICs) (Figure 1C), suggesting that these proteins form HTT-associated multiprotein complexes within the mitochondria. Similar results were observed when we immunoprecipitated HTT from mouse brain tissue (3-month-old C57BL/6 wild-type mice) the same proteins appeared in HTT ICs (Figure 1D). Since TFAM was present in the HTT ICs and forms a complex with the proteins IP’d using anti-HTT antibody, we performed IP using anti-TFAM antibody to pull down TFAM-associated proteins from the MEs from mouse brain to further validate our findings. IP of TFAM from MEs with anti-TFAM antibody showed identical proteins to those IP’d using the anti-HTT antibody (Figure 1E). IP of POLRMT with anti-POLRMT antibody from the MEs isolated from wildtype mouse brain tissue confirmed the presence of same protein interactions as in the HTT ICs (Figure 1F). Finally, IP of mitochondrial PNKP from wildtype mouse brain MEs confirmed the presence of the same proteins in the PNKP ICs as in HTT ICs (Figure 1G). Importantly, we detected Cockayne syndrome protein B (CSB) in all ICs (HTT, TFAM, POLRMT, and PNKP ICs; Figures 1B to 1G), which is known to regulate mtDNA repair [28, 29]. These findings suggest that HTT forms a complex with POLRMT, POLGA, TFAM, CSB and PNKP within mitochondria of neuron-like cells. To ensure interaction specificity, we tested for TOM20 (an outer mitochondrial membrane protein) in the ICs. No TOM20 was detected (Figures 1B to 1G), suggesting specificity of these protein interactions.

### HTT-TFAM interactions increase upon mtDNA damage

While the biochemical experiments demonstrate HTT forms a complex with the mitochondrias-pecific proteins that are known to be important in mtDNA transcription and repair, it does not show whether HTT interacts with these proteins *in situ* and whether these interactions are altered when damage is induced in mtDNA. To test this, we performed proximity ligation assays (PLAs) to assess *in situ* protein interactions[21], in neuronal SH-SY5Y cells before and after inducing mtDNA damage. We treated cells with glucose oxidase (GO; 300 ng/mL for 30 min), an oxidizing agent that generates hydrogen peroxide *in situ* and preferentially induces mtDNA damage[30]. We first confirmed mtDNA damage using long-amplicon quantitative polymerase chain reaction (LA-QPCR) analysis[21, 31]. As expected, LA-QPCR analysis showed reduced PCR amplification efficiency for mtDNA (8.9 kb segment) in GO-treated versus control cells, confirming substantial mtDNA damage (Figures 2A and 2B). Consistent with biochemical findings, notable fluorescent signals emerged when we performed PLA on SH-SY5Y cells using anti-TFAM and anti-HTT antibodies (green, arrows). These PLA signals strongly overlapped with mitochondria (marked by MitoTracker Red; arrows), indicating that HTT and TFAM interact within mitochondria in living cells (Figure 2C; arrows). Strikingly, the fluorescence reconstitution signals increased 3 to 4-fold and overlapped with MitoTracker Red following GO-induced DNA damage (Figures 2D; arrows, and 2E), suggesting an increased HTT-TFAM association in response to increased mtDNA damage.

**Figure 2:**
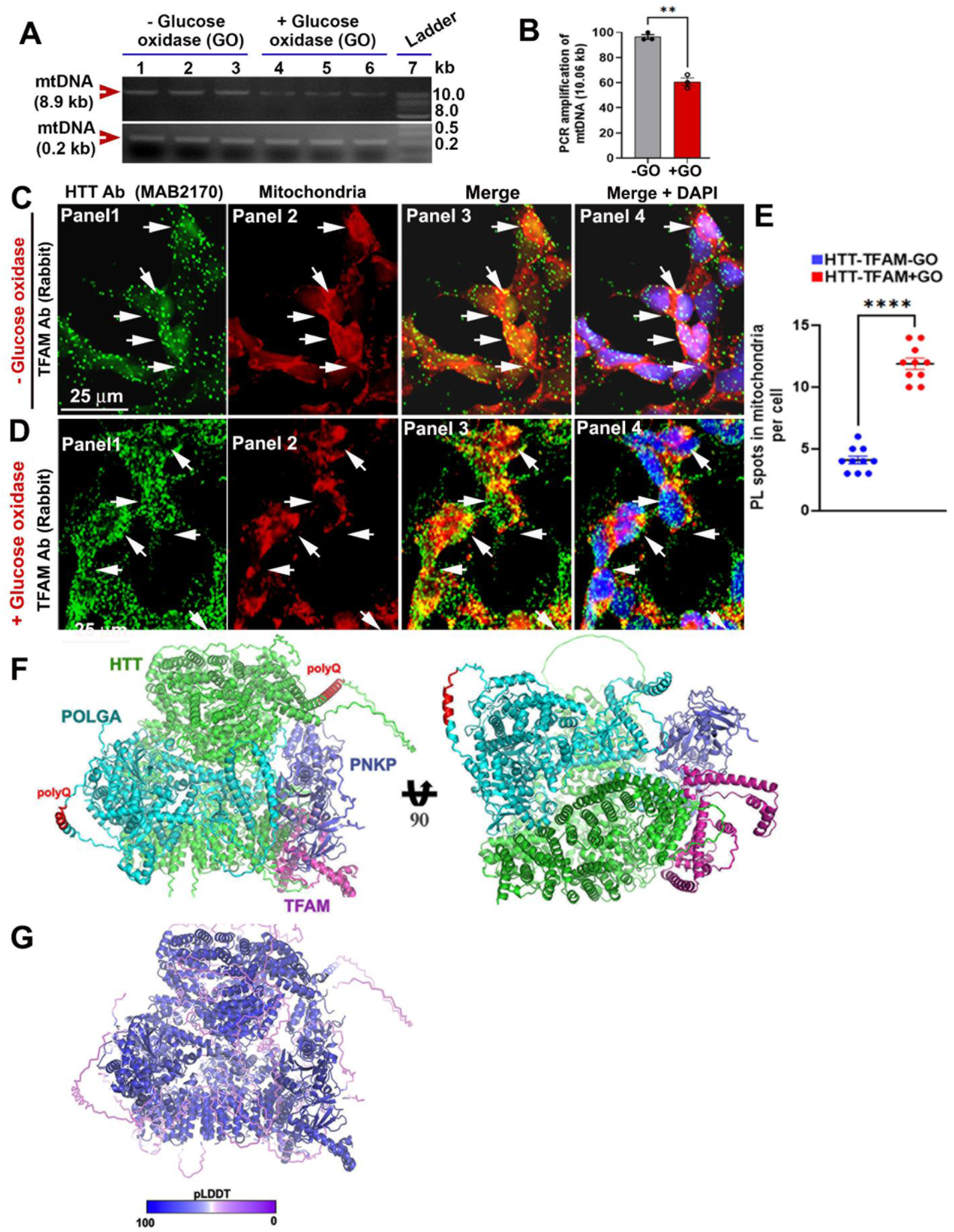
Association of HTT with the mitochondrial transcription complex is increased in response to oxidative stress *in vitro*. **(A).** SH-SY5Y cells treated with glucose oxidase (GO; 300 ng/mL) for 30 minutes, total DNA isolated from untreated control (lanes 1 to 3) and GO-treated (lanes 4 to 6) cells, and 8.9 kb and 0.2 kb segments of mtDNA PCR amplified, and the PCR products electrophoresed on agarose gels, and the DNA bands quantified. The amplification efficiencies of the 8.9 kb and 0.2 kb segments of mtDNA are shown by arrowheads. Lane 7: 1-kb DNA ladder. **(B).** Relative PCR amplification efficiencies of mtDNA from control and GO-treated SH-SY5Y cells as described in Figure 2A. Data represent mean ± SD; **p<0.01. Proximity ligation assay (PLA) performed to assess possible interaction of HTT with mitochondrial transcription factor TFAM or mitochondrial RNA polymerase (POLRMT) in mitochondria in SH-SY5Y cells. Mitochondria stained with Mito Tracker Red (panels 2), and generation of green fluorescence (panels 1) indicate representative positive protein-protein interactions (arrows in panels 3 and 4 show positive PLA signals in yellow inside mitochondria). Nuclei stained with DAPI (panels 4). PLA with mouse monoclonal anti-HTT antibody (MAB 2170; EMD Millipore) and anti-TFAM Ab (ab176558; Abcam) or anti-POLRMT rabbit polyclonal Ab (GTX105137; Genetex) performed to assess interaction of HTT with TFAM or POLRMT. Cells then treated with glucose oxidase (GO: 300 ng/mL) for 30 minutes to induce the mtDNA damage. **(C).** PLA performed on SH-SY5Y cells without GO treatment to assess association/interactions of endogenous HTT with TFAM within mitochondria. Nuclei are stained with DAPI in C, D, F, and G. **(D).** SH-SY5Y cells treated with GO (300 ng/mL) for 30 minutes to induce mtDNA damage, and PLA performed to assess possible interactions of endogenous HTT with TFAM. **(E).** Increased PLA signals showing interaction of TFAM and HTT within mitochondria in GO-treated (+GO) cells compared to control (-GO) cells. Data represent mean ± SD; ****p<0.0001. **(F).** HTT-POLGA-TFAM-PNKP computational model based upon AlphaFold 3. **(G).** pLDDT model confidence maps onto the predicted model. Blue denotes higher confidence, and purple denotes lower confidence.

In addition, a HTT -POLRMT interaction was observed within mitochondria, and this interaction was significantly increased (increased PLA signal overlapping with MitoTracker Red signal) upon GO-induced mtDNA damage (Supplementary figures 1A to 1C). For additional validation, we IP’d FLAG-wtHTT from the MEs of human HEK293 cells co-expressing FLAG-wtHTT and Myc-tagged POLRMT, POLGA, PNKP or TFAM using anti-FLAG antibody. WB analyses of FLAG ICs confirmed the presence of POLRMT, POLGA, PNKP and TFAM in the FLAG IC (Supplementary figures 1D and 1E), suggesting wtHTT directly interacts with POLGA, POLRMT, PNKP and mitochondrial transcription factor TFAM. To assess whether HTT interacts with endogenous POLRMT, POLGA, PNKP or TFAM, we expressed FLAG-HTT-Q24 in SH-SY5Y cells, isolated the MEs, and IP’d the exogenous FLAG-tagged HTT from the MEs with anti-FLAG antibody (F3165; Millipore-Sigma). Western blotting analysis of FLAG ICs revealed the presence of endogenous POLRMT, POLGA, and PNKP in the FLAG ICs (Supplementary figure 1F), further supporting the interpretation that HTT interacts with endogenous mitochondria-specific proteins e.g., POLRMT and PNKP, and forms a multifactorial complex within cellular mitochondria. We next performed bimolecular fluorescence complementation (BIFC) to test whether HTT interacts with POLRMT or POLGA within the mitochondria. A substantial reconstitution of green fluorescence within the mitochondria suggests interactions of HTT with mitochondrial proteins POLGA and POLRMT (Supplementary figures 2G and 2H; arrows).

In addition, significant PLA signals indicated a strong association between PNKP and TFAM. This association was increased upon GO treatment, showing enhanced interaction between TFAM and PNKP in the mitochondria in response to increased mtDNA damage (Supplementary figures 2A to 2C). PLA also showed significantly increased recruitment of CSB, a key protein essential for initiating transcription-coupled DNA repair[32–34] into the HTT-POLRMT complex in response to mtDNA damage (Supplementary figure 2D to 2F). These data demonstrate that wtHTT forms a multifactorial transcription-linked DNA repair complex with important components of the mitochondrial transcription and replication machinery and it associates with mtDNA. Importantly, the recruitment of HTT to mtDNA is markedly enhanced upon induction of mtDNA damage. We next performed bimolecular fluorescence complementation (BIFC) to test whether HTT interacts with POLRMT or POLGA within the mitochondria. A substantial reconstitution of green fluorescence within the mitochondria suggests substantial interactions of HTT with mitochondrial proteins POLGA and POLRMT (Supplementary figures 2G and 2H; arrows).

Despite HTT’s large size making computational modeling challenging, we reasoned that AlphaFold 3 (AF3) a computational structure prediction could provide an independent assessment of the potential for structural interactions within the mt DNA repair complex[35]. The AF3 computational structure supported HTT-mediated protein assembly with POLGA, PNKP, and TFAM (Figure 2F). Notably, AF3 places HTT in the core of this complex with PNKP being tucked between HTT, POLGA and TFAM (Figure 2F). Finally, to test human relevance of these findings, we examined *post-mortem* human brain tissue (cortex), where substantial PLA signals were detected between wtHTT and POLRMT or POLGA (Supplementary figures 3A and 3B; arrows). These results support that wtHTT interacts with POLRMT and POLGA within human neuronal mitochondria. These data show that wtHTT and PNKP are present within the mitochondria and form a macromolecular complex with POLRMT, DNA polymerase γ and associated mitochondrial transcription factors.

Next, we reasoned that if wtHTT and its interacting partner PNKP[21] were to form a mtDNA repair complex, HTT and PNKP could be associated with mtDNA *in vivo*. Chromatin immunoprecipitation (ChIP) analysis was performed to examine the possible interactions between HTT and PNKP with mtDNA using mouse brain tissue (Cortex). ChIP analyses revealed notable interactions between endogenous HTT and mtDNA in mouse brain tissue (Figure 3A), suggesting that the native endogenous HTT physically interacts with mtDNA under physiological conditions *in vivo*. Furthermore, ChIP analysis also showed notable *in vivo* association/interaction of endogenous PNKP with mtDNA using tissue lysate derived from wildtype mouse brain under identical experimental conditions (Figure 3B). These data strengthen the interpretation that HTT and PNKP are present within the mitochondria and physically interact with mtDNA and likely play important roles in mtDNA damage repair[27].

**Figure 3:**
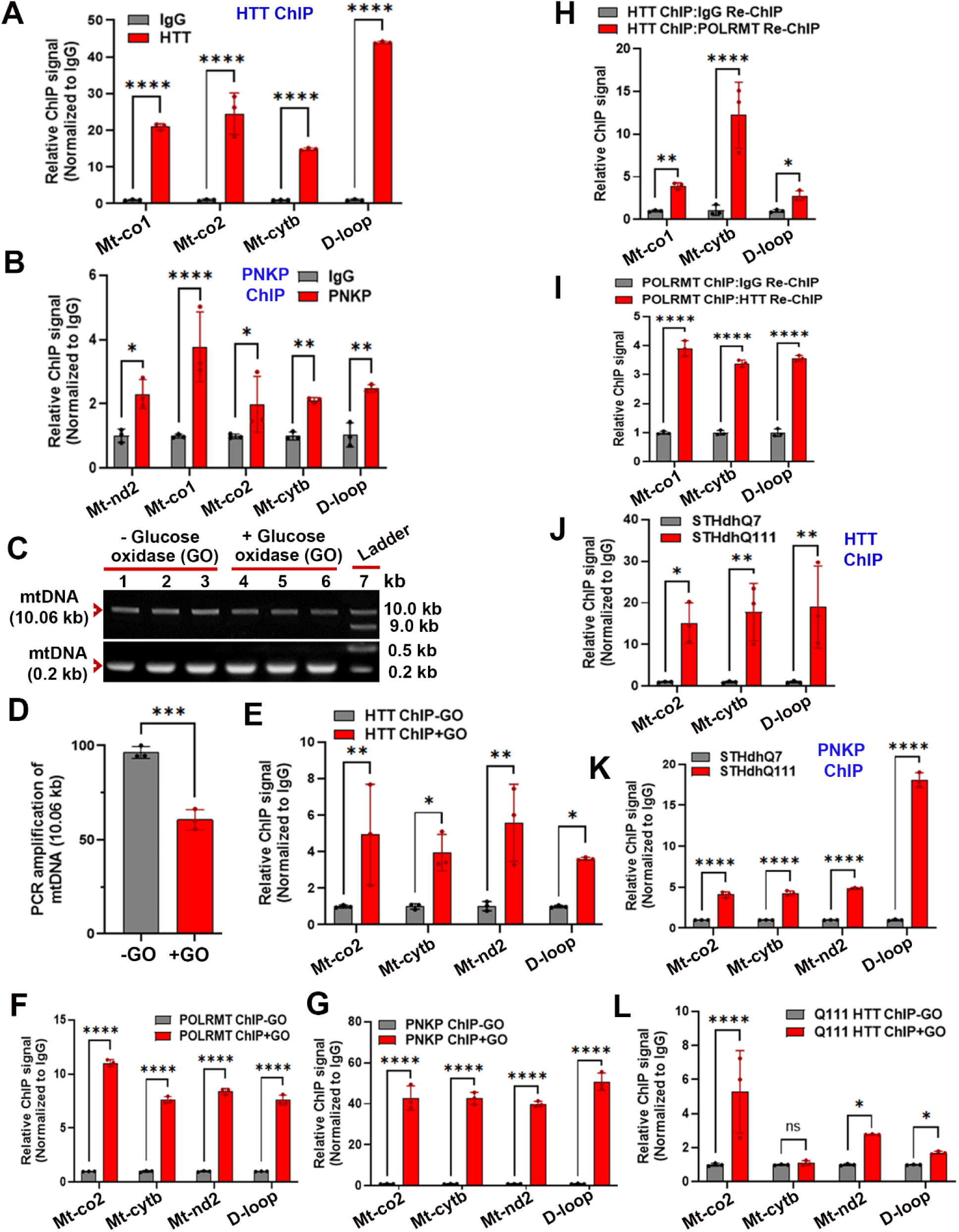
HTT and PNKP interact with mtDNA, and these interactions are increased upon oxidative stress-induced mtDNA damage. **(A).** Chromatin immunoprecipitation (ChIP) analysis performed to assess the possible interaction of endogenous native HTT with mtDNA in wildtype, male C57BL/6 mouse cortex (3-month-old). ChIP showing relative association of wtHTT with various segments of mtDNA sequences e.g., cytochrome c oxidase 1 and 2 (Mt-co1 and Mt-co2 respectively), mitochondrial cytochrome b (Mt-cytb), and the mitochondrial D-loop sequences in wildtype (3-months-old) male C57BL/6 mouse brain tissue. The relative ChIP values were measured after normalization to control IgG in each case. Three biological replicates, and three technical replicates were used for this measurement and for B, E, F, G, H, I, and J. Data represent mean ± SD: ****p<0.0001. **(B).** ChIP showing *in vivo* association of endogenous PNKP with various segments of mtDNA (Mt-nd2, Mt-co1, Mt-co2, Mt-cytb and D-loop) sequences in 3-months-old C57BL/6 male mouse brain tissue. Relative ChIP values normalized to control IgG. Data represents mean ± SD. *p<0.05; **p<0.01; ****p<0.0001. **(C).** STHdhQ7 striatal cells treated with glucose oxidase (GO; 300 ng/mL) for 30 minutes. After GO treatment, cells harvested and total DNA isolated from the untreated control (lanes 1 to 3) and GO-treated (lanes 4 to 6) cells and 10.06 kb and 0.2 kb segments of mouse mtDNA PCR amplified, and the PCR products electrophoresed on agarose gels, and the intensity of the DNA bands quantified. The amplification efficiencies of the 10.6 kb and 0.2 kb segments of mouse mtDNA are shown by arrowheads. Lane 7: 1-kb DNA ladder. **(D).** Relative PCR amplification efficiencies of mtDNA from control and GO-treated STHdhQ7 cells as described in figure 3C. Data represent mean ± SD; ***p<0.001. **(E).** Striatal wildtype STHdhQ7 cells treated with GO (300 ng/mL) for 30 minutes, cells harvested, and ChIP performed on GO-treated and untreated control cells to determine the relative interaction of endogenous native HTT with mtDNA before and after inducing mtDNA damages with GO. ChIP analysis showing relative association of HTT with Mt-co2, Mt-cytb, Mt-nd2 and D-loop sequences in GO-treated vs. untreated cells. The relative ChIP values measured after normalization to control IgG in each case. Data represents mean ± SD; *p<0.05; **p<0.01. **(F).** Striatal STHdhQ7 cells treated with GO (300 ng/mL) for 30 minutes, cells harvested, and ChIP performed on GO-treated and untreated cells to determine the relative interaction of mitochondrial RNA polymerase POLRMT with mtDNA in presence or absence of mtDNA damages. ChIP showing relative association of POLRMT with Mt-co2, Mt-cytb, Mt-nd2 and D-loop sequences in GO-treated vs. untreated cells. The relative ChIP values measured after normalization to control IgG in each case. Data represents mean ± SD; ****p<0.0001. **(G).** STHdhQ7 cells were treated with GO (300 ng/mL) for 30 minutes, cells were harvested, and ChIP performed on GO-treated and untreated cells to determine the relative interaction of PNKP with mtDNA in presence or absence of mtDNA damages. ChIP showing relative association of PNKP with Mt-co2, Mt-cytb, Mt-nd2 and D-loop sequences in GO-treated vs. untreated control cells. The relative ChIP values measured after normalization to control IgG in each case. Data represent mean ± SD; ****p<0.0001. **(H).** ChIP-re-ChIP analysis performed to assess co-occupancy of HTT and POLRMT with mtDNA in wildtype C57BL/6 mouse brain tissue. Putative HTT-mtDNA complexes isolated, and the complexes immunoprecipitated with an anti-HTT mouse monoclonal Ab (MAB2170: Millipore-Sigma), followed by a second IP from the HTT ICs with an anti-POLRMT rabbit polyclonal Ab (GTX105137; Genetex) or anti-IgG Abs. DNA isolated from the final POLRMT ICs and IgG IC, and various mtDNA segments encoding Mt-co1, Mt-cytb, and D-loop sequences amplified by quantitative PCR. Relative ChIP values measured after normalization to control IgG. Data represents mean ± SD; *p<0.05; **p<0.01; ****p<0.0001. **(I).** The reverse method of ChIP-re-ChIP analysis performed. Putative POLRMT-mtDNA complexes sequentially immunoprecipitated using an anti-POLRMT rabbit polyclonal Ab (GTX105137; Genetex) followed by anti-HTT mouse Ab (MAB2170; Millipore-Sigma), and respective control anti-IgG Abs. The presence of mtDNA segments in the final HTT IC or IgG IC detected by quantitative PCR as described in F. Data represent mean ± SD; ****p<0.0001. **(J).** ChIP performed to assess possible interaction of the wildtype HTT (wtHTT) and mutant HTT (mHTT) with mtDNA in wildtype STHdhQ7 and mutant STHdhQ111 cells expressing wtHTT or mHTT respectively. ChIP showing relative association of wtHTT and mHTT with various segments of mtDNA sequences e.g., Mt-co2, Mt-cytb, and D-loop sequences in STHdhQ7 or STHdhQ111 cells. The relative ChIP values measured after normalization to control IgG in each case. Three biological replicates each with three technical replicates used for this measurement. Data represents mean ± SD; *p<0.05; **p<0.01. **(K).** ChIP performed to assess possible interaction of PNKP with mtDNA of STHdhQ7 and STHdhQ111 cells. ChIP showing relative association of PNKP with various segments of mtDNA sequences e.g., Mt-co2, Mt-cytb, Mt-nd2, and mitochondrial D-loop sequences in STHdhQ7 or STHdhQ111 cells. The relative ChIP values measured after normalization to control IgG in each case. Three biological replicates each with three technical replicates used for this measurement. Data represent mean ± SD; ****p<0.0001. **(L).** Homozygous mutant STHdhQ111 cells treated with GO (300 ng/mL) for 30 minutes, cells harvested, and ChIP performed on GO-treated and control untreated cells to determine the relative interaction of mHTT with mtDNA in presence or absence of mtDNA damages. ChIP showing relative association of HTT with Mt-co2, Mt-cytb, Mt-nd2 and D-loop sequences in GO-treated vs. untreated STHdhQ111 cells. The relative ChIP values measured after normalization to control IgG in each case. Data represents mean ± SD; *p<0.05; ****p<0.0001; ns = not significant.

Having established that both wtHTT and its interacting DNA repair enzyme PNKP[21] can bind to the mtDNA *in vivo*, and interact with other proteins required for mitochondrial transcription, we tested whether wtHTT and associated proteins directly participate in mtDNA repair. We reasoned that if wtHTT and its interacting proteins are required for mtDNA repair, their association with mtDNA may increase upon increased mtDNA damage. To test this, we induced mtDNA damage in striatal STHdhQ7 cells expressing wtHTT[36] and assessed the interactions between wtHTT and PNKP with mtDNA. We treated the STHdhQ7 cells with GO (300 ng/mL for 30 min) to induce damage in mtDNA followed by LA-QPCR analyses [21, 31] to assess mtDNA damage before and after the GO treatment. As expected, GO treatment induced substantial mtDNA damage as revealed by LA-QPCR analysis (Figures 3C and 3D). Also, ChIP analysis showed a dramatic increase in the association of endogenous wtHTT with mtDNA in GO-treated STHdhQ7 cells (Figure 3E), suggesting an increased recruitment/association of HTT with mtDNA in response to increased mtDNA damage. As a positive control, we assessed POLRMT-mtDNA interactions in STHdhQ7 cells before and after GO-induced DNA damage. As expected, ChIP analysis confirmed significantly increased POLRMT-mtDNA association upon GO treatment (Figure 3F). Similarly, the ChIP analysis shows that the interaction/association of PNKP with mtDNA is increased by several fold in GO-treated cells compared with control (Figure 3G).

To confirm these protein-mtDNA interactions using another independent method, we performed sequential ChIP (ChIP-re-ChIP) analysis[37]. First, we immunoprecipitated putative HTT-bound mtDNA fragments with an anti-HTT antibody, followed by IP of the wtHTT immunocomplexes (ICs) with an anti-POLRMT or with an anti-IgG antibody. The quantitative PCR (QPCR) analysis of DNA isolated from the final POLRMT ICs showed a robust amplification of various mtDNA segments (Figure 3H). We also performed the converse experiments where in the protein-DNA interactors were first IP’d with an anti-POLRMT antibody, then subjected to a second IP of the POLRMT ICs with an anti-HTT antibody or with an IgG antibody followed by QPCR analysis of the DNA fragments isolated from the final HTT and IgG ICs. A significant amplification of the same mtDNA segments from DNA isolated from the final HTT ICs was observed (Figure 3I).

Next, we investigated whether the polyQ expansion in HTT alters the mtDNA-mHTT interaction by comparing relative interactions between mtDNA and mHTT in STHdhQ111 cells (expressing mHTT) and with HTT and mtDNA in control wildtype STHdhQ7 cells (expressing HTT) verses HTT-mtDNA interaction in control STHdhQ7 cells (expressing mHTT) [36]. ChIP analysis revealed significant interactions between HTT and mtDNA in both controls and in mutant cells (Figure 3J). Intriguingly, a significantly higher (12-15-fold; p<0.001) association/occupancy of mHTT with mtDNA was observed as compared to the interaction between HTT and mtDNA (Figure 3J), suggesting that mHTT may associate/bind with the mtDNA with a higher affinity and therefore the occupancy of mHTT with mtDNA is higher in mutant cells. Importantly, ChIP analysis also showed a significantly higher association between PNKP and mtDNA in mutant STHdhQ111 cells compared with the wildtype STHdhQ7 cells (Figure 3K), indicating that PNKP may be recruited more to mtDNA in mutant cells due to the presence of higher mtDNA damage in mutant STHdhQ111 cells. Further, ChIP analysis revealed a significant interaction of mHTT with mtDNA in STHdhQ111 cells that constitutively express mHTT, and this protein-mtDNA interaction is markedly increased when mtDNA damage was induced by GO (Figure 3L).

Taken together, these results strongly suggest that HTT forms a mtDNA repair complex *in vivo* and perhaps, a multiprotein transcription-linked DNA repair complex together with mitochondrial transcription factors (although the latter requires further evaluation).

### Wildtype HTT facilitates mtDNA repair and maintenance of mitochondrial genome integrity and function

The data above suggest that wtHTT may participate in mtDNA repair, therefore we tested whether wtHTT could protect mitochondria by stimulating mtDNA repair and maintaining mitochondrial genome integrity. We assessed the mtDNA repair activity in wtHTT-depleted (HTT-RNAi) SH-SY5Y cells by measuring the PNKP activity in the purified MEs. Depleting wtHTT levels in SH-SY5Y cells significantly reduced mitochondria-specific PNKP activity, but the steady-state levels of PNKP, POLRMT and TFAM remain unchanged (Figures 4A to 4C), suggesting that HTT may regulate the enzymatic activity of mitochondrial PNKP. This observation mirrors our previous finding that HTT regulates PNKP activity in the nucleus[21]. Importantly, depleting HTT also resulted in increased accumulation of mtDNA lesions compared to control-RNAi-expressing cells (Figures 4D and 4E). Further, HTT-overexpressing (HTT-OE) SH-SY5Y cells showed a marked increase in mtDNA repair activity (Figures 4F to 4H) and improved mtDNA integrity, demonstrated by LA-QPCR analysis (Figures 4I and 4J). These data suggest that cellular wtHTT levels regulate both PNKP activity and the efficacy of mtDNA repair. To test whether increased HTT levels might provide added protection through enhanced DNA repair capability, we exposed SH-SY5Y cells to pro-degenerative stimuli. Indeed, upon treatment of HTT-OE SH-SY5Y cells with GO, they showed higher resistance to mtDNA damage compared to HTT-KD SH-SY5Y and control cells (Figures 4K and 4L). Cellular toxicity was measured using an MTT assay kit. As expected, the HTT-KD cells showed higher cell toxicity compared with HTT-OE cells (Figure 4M). We next measured mitochondrial membrane potentials in control cells, HTT-KD and HTT-OE cells to test whether HTT levels also determine mitochondrial function. To test this, we treated control SH-SY5Y, HTT-KD, and HTT-OE cells with the complex II inhibitor 3-NP in the presence of JC-10 dye. In polarized cells, JC-10 dye predominantly concentrates in the mitochondrial matrix where it forms red/orange aggregates. In contrast, in depolarized cells, JC-10 diffuses out of mitochondria and exists primarily as a monomer, which stains cells green. While HTT-KD cells showed significantly lower mitochondrial membrane potential (Green fluorescence), HTT-OE cells showed significantly higher mitochondrial membrane potential (red/orange fluorescence) compared with control cells (Figure 4N), indicating the preservation of mitochondrial function. Finally, depleting other key components of the HTT-interacting mitochondrial proteins (e.g., POLRMT or PNKP) also impaired the cellular ability to efficaciously repair mtDNA damage (Supplementary figure 4), indicating that these proteins are important for effective mtDNA repair. Together, these data show that HTT protects both mtDNA as well as mitochondrial function.

**Figure 4.**
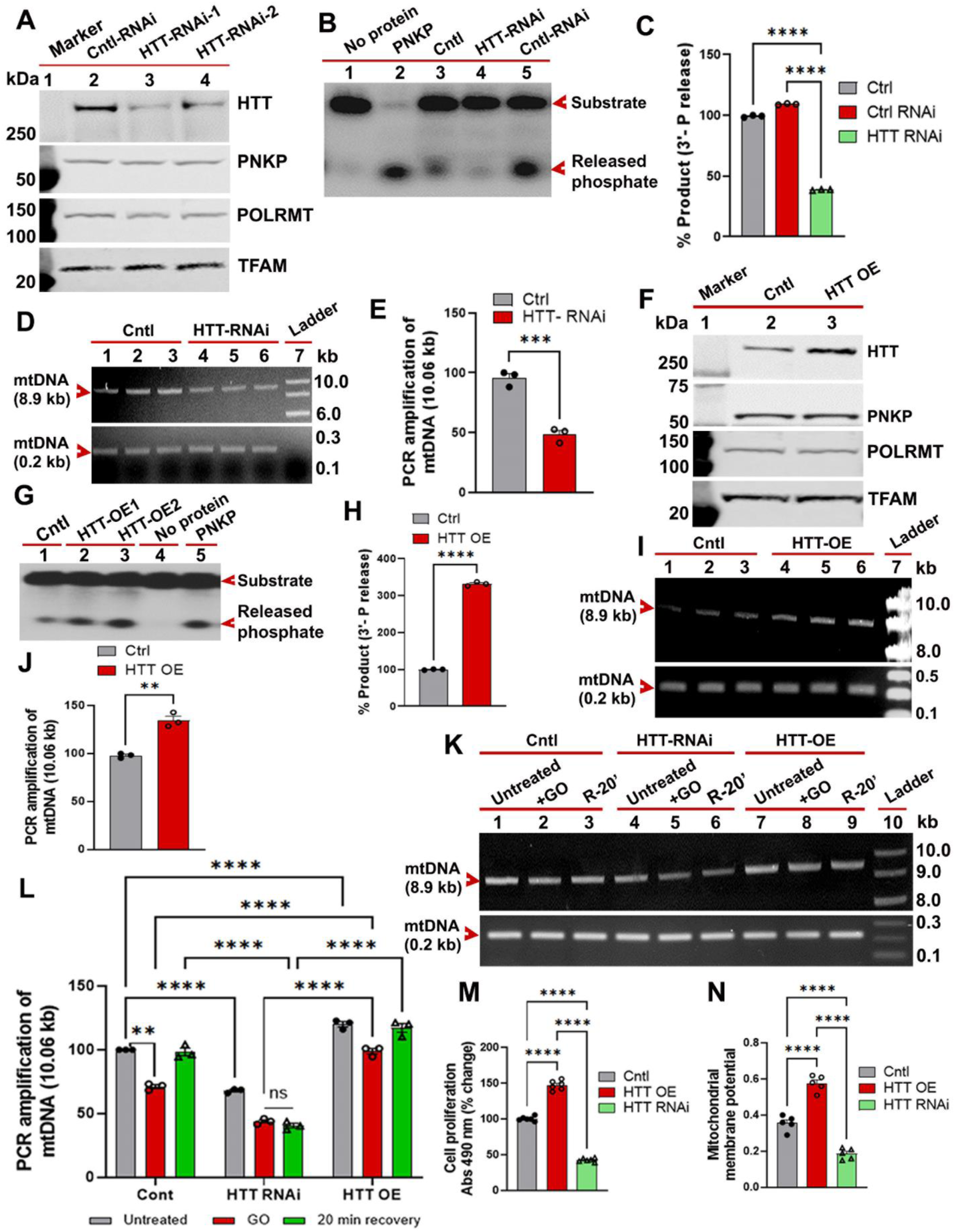
Wildtype HTT (wtHTT) stimulates mtDNA damage repair to maintain mitochondrial genome integrity and mitochondrial function. **(A).** Total proteins extracted from SH-SY5Y cells, and SH-SY5Y cells expressing control-RNAi (Cntl-RNAi; lane 2) or two different HTT-RNAi (HTT-RNAi-1 and HTT-RNAi-2; lanes 3 and 4), and the protein fractions analyzed by WBs to detect the expression levels of HTT, PNKP, POLRMT and TFAM; Lane 1: protein molecular weight marker in kDa; Mitochondrial transcription factor TFAM used as loading control. **(B).** The 3’-phosphatase activities of PNKP in the MEs (250 ng each) from wildtype control SH-SY5Y cells (Cntl: lane 3), SH-SY5Y cells expressing HTT-RNAi (HTT-RNAi: lane 4) or Control-RNAi (Cntl-RNAi-1; lane 5) determined by adding MEs to the radiolabeled (32-P) DNA substrate (blue arrowhead) and measuring the release of the radio-labelled phosphate (red arrowhead) from the substrate. Lane 1: No protein extract added to the substrate as negative control; Lane 2: Purified PNKP protein (25 femtomoles) added to the substrate as a positive control. **(C).** Relative 3’-phosphatase activities of PNKP (in terms of % product) in wildtype SH-SY5Y cells (Cntl), and SH-SY5Y cells either expressing control-RNAi (Cntl-RNAi) or HTT-RNAi. Data represents mean ± SD. Three biological replicates and three technical replicates used for this measurement and in E, H, J, L, M, and N. Data represent mean ± SD: ****p<0.0001. **(D).** Total DNA isolated from control SH-SY5Y cells (Cntl; lanes 1-3), and SH-SY5Y cells expressing HTT-RNAi (lanes 4-6), 8.9 kb and 0.21 kb segments of mtDNA PCR amplified, and the PCR products analyzed on agarose gels (arrowheads). Lane 7: 1-kb DNA ladder. **(E).** Relative PCR amplification of an 8.9 kb segment of mtDNA isolated from the control SH-SY5Y cells (Cntl), and SH-SY5Y cells constitutively expressing HTT-RNAi. Data represents mean ± SD: ***p<0.001. **(F).** Control SH-SY5Y cells (Cntl) or SH-SY5Y cells expressing exogenous full-length wtHTT-Q23 (HTT-OE-1) harvested, and mitochondrial protein extracts isolated, and the MEs analyzed by WBs to detect mitochondrial HTT, PNKP, POLRMT and TFAM levels; Lane 1: protein molecular weight marker; TFAM used as loading control. **(G).** The 3’-phosphatase activities of PNKP in MEs (250 ng each) from wildtype control SH-SY5Y cells (Cntl: lane 1), and SH-SY5Y cells over-expressing wtHTT (HTT-OE-1 or HTT-OE-2: lanes 2 and 3) determined by adding MEs to the radiolabeled DNA substrate (red arrowhead) and measuring the phosphate release (red arrowhead) from the DNA substrate. Lane 4: No protein extract added to the substrate as negative control; Lane 5: Purified PNKP protein (25 femtomoles) added to the substrate as a positive control. **(H).** Relative 3’-phosphatase activities of PNKP (in terms of % product) in the control SH-SY5Y cells (Cntl), and in SH-SY5Y cells overexpressing exogenous wtHTT-Q23 (HTT-OE). Data represent mean ± SD; ****p<0.0001. **(I).** DNA isolated from control SH-SY5Y cells (Cntl; lanes 1 to 3), and HTT-OE cells (lanes 4 to 6), 8.9 kb and 0.21 kb sections of mtDNA PCR amplified, and the PCR products analyzed on agarose gels and stained with Ethidium bromide (EtBr) (arrowheads). Lane 7: 1 kb DNA ladder. **(J).** Relative mtDNA integrity in control (Cntl) SH-SY5Y cells, and SH-SY5Y cells overexpressing wtHTT (HTT-OE); Data represent mean ± SD; **p<0.01. **(K).** Control SH-SY5Y cells (Cntl; lanes 1 to 3), SH-SY5Y cells expressing HTT-RNAi (lanes 4 to 6), or SH-SY5Y cells overexpressing wtHTT (HTT-OE; lanes 7 to 9) treated with glucose oxidase (GO; 300 ng/mL; lanes 2, 3, 5, 6, 8 and 9) for 60 minutes. Cells introduced with fresh medium for the repair of mtDNA damage, and cells harvested after 20 minutes of repair (R-20’; lanes 3, 6 and 9), total DNA isolated from each cell pellet, and 8.9 kb and 0.2 kb segments of mtDNA PCR amplified, and the PCR products analyzed on agarose gels to determine the relative PCR amplification of mtDNA (arrowheads). Lane 10: 1-kb DNA ladder. **(L).** Relative levels of mtDNA damage in control SH-SY5Y cells (Cntl), HTT-RNAi cells, and HTT-OE cells, after treating cells with GO (300 ng/mL) for 60 minutes, and 20 minutes after GO removal (Recovery-20’); Data represent mean ± SD; **p<0.01; ****p<0.0001; ns = not significant. **(M).** Cell viabilities (MTT assay) measured in control SH-SY5Y cells (Cntl), SH-SY5Y cells expressing exogenous wtHTT-Q23 (HTT-OE), or HTT-RNAi; Data represent mean ± SD; ****p<0.0001. **(N).** Mitochondrial membrane potential measured with JC-10 dye in control SH-SY5Y cells (Cntl), HTT-RNAi cells and HTT-OE cells; Data represent mean ± SD; ****p<0.0001.

### Mutant HTT impairs efficacy of mtDNA damage repair

We previously demonstrated that mutant HTT (mHTT) impairs nuclear DNA repair by disrupting the catalytic activity of PNKP[21], which led us to investigate whether mHTT had an effect also in mitochondria. We first asked whether mHTT interferes with mitochondrial PNKP activity and mtDNA repair in an immortalized striatal cell model of HD[36]. We isolated mitochondria from mutant STHdhQ111 and control wildtype STHdhQ7 cells and used purified MEs from these cells to measure the 3’-phosphatse activity of PNKP and the efficacy of mtDNA repair. MEs from the STHdhQ111 cells showed significantly lower (∼70-75%) PNKP activity compared with STHdhQ7 cells, while PNKP protein levels remained unaltered between these cells (Figures 5A and 5B). Since STHdhQ111 cells showed reduced PNKP activity compared with STHdhQ7 cells, we tested whether mutant STHdhQ111 cells have higher mtDNA damage, and whether increasing PNKP activity could restore efficacy of mtDNA repair. To test this, we overexpressed PNKP in STHdhQ111 cells (Figure 5C; lanes 7-9 vs. lanes 1-3) and assessed mtDNA repair efficacy in mutant cells and mutant cells expressing human PNKP using LA-QPCR[21, 31]. We observed significantly higher mtDNA damage in mutant STHdhQ111 cells compared with control STHdhQ7 cells (Figures 5D; lanes 4-6 vs. lanes 1-3, and 5E). Importantly, mtDNA integrity was significantly restored when PNKP levels were elevated in mutant STHdhQ111 cells (Figures 5D; lanes 7-9 vs. lanes 4-6, and 5E). Finally, increasing PNKP levels in STHdhQ111 cells also resulted a significant restoration of mitochondrial membrane potential and reduced cell toxicity (Figures 5F and 5G), indicating that mHTT disrupts mitochondrial PNKP activity, and restoring PNKP activity in mutant cells increased mtDNA repair efficacy and restored mitochondrial function with a significant reduction in cell toxicity.

**Figure 5:**
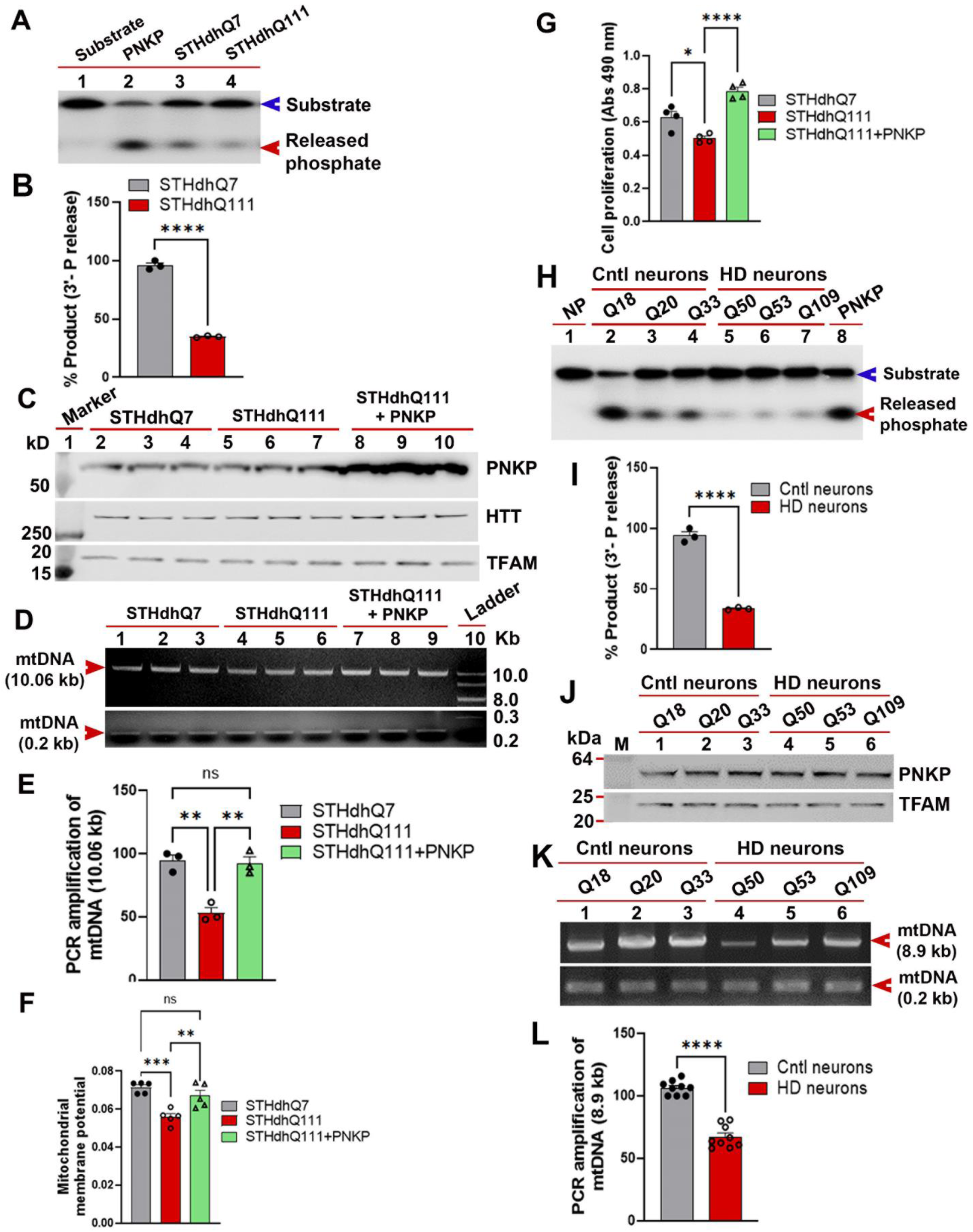
Mutant HTT (mHTT) dramatically reduces mitochondrial PNKP activity, resulting in impaired mtDNA repair, and accumulation of mtDNA damage *in situ*. **(A).** PNKP’s 3’-phosphatase activities in MEs of control STHdhQ7 (lane 3) and mutant STHdhQ111 cells (l and 4) measured by the 3’-phosphate release (red arrowhead) from the DNA substrate (blue arrowhead). Lane 1: No protein extract added to the DNA substrate as negative control; Lane 2: Purified PNKP (25 fmoles) added to the substrate as positive control. **(B).** Relative 3’-phosphatase activities of PNKP (in terms of % product) in mutant STHdhQ111 and control STHdhQ7 cells. Three biological and three technical replicates used for calculating PNKP activity. Data represents mean ± SD; ****p<0.0001. **(C).** Proteins were isolated from STHdhQ7 (lanes 2 to 4), STHdhQ111 (lanes 5 to 7) and STHdhQ111 cells expressing exogenous human PNKP (lanes 8 to 10), analyzed by WBs to detect PNKP and HTT levels; Lane 1: Protein molecular weight marker in kDa; Mitochondrial transcription factor TFAM used as loading control. **(D).** STHdhQ7 (lanes 1 to 3), STHdhQ111 (lanes 4 to 6) and STHdhQ111 cells expressing exogenous human PNKP (lanes 7 to 9) harvested, total DNA isolated and 10.06-and 0.2-kb sections of mouse mtDNA segments PCR amplified; and the products analyzed on agarose gels (red arrowheads). Lane 10: 1-kb DNA ladder. **(E).** Relative PCR amplification of mtDNA in STHdhQ7, STHdhQ111, and STHdhQ111 cells expressing exogenous PNKP. Three biological and three technical replicates were used for the calculation. Data represent mean ± SD; **p<0.01; ns = not significant. **(F).** Control STHdhQ7 cells, and mutant STHdhQ111 cells and STHdhQ111 cells overexpressing exogenous PNKP treated with JC-10 dye and mitochondrial membrane potential measured; Three biological replicates and three chemical replicates were used for this measurement; Data represents mean ± SD; **p<0.01; **p<0.001; ns = not significant. **(G).** Cell viabilities of STHdhQ7, STHdhQ111 and STHdhQ111 cells expressing exogenous PNKP measured; Data represent mean ± SD; *p<0.05; ****p<0.0001. **(H).** The 3’-phosphatase activities of PNKP in the MEs (250 ng each) of control (lanes 2, 3 and 4, differentiation of wtHTT-Q18, Q20 and Q33 iPSCs), and HD striatal neurons (lanes 5, 6 and 7, differentiation of mHTT-Q50, Q53 and Q109 iPSCs) determined by adding the MEs to the DNA substrate and measuring the 3’-phosphate release (red arrowhead) from the DNA substrate (blue arrowhead). Lane 1: No protein extract (NP) added to the DNA substrate as negative control; Lane 8: Purified PNKP (25 femtomoles) added to the DNA substrate as positive control. **(I).** Relative 3’-phosphatase activities (in terms of % product) of mitochondrial PNKP in control (Q18, Q20 and Q33) vs. HD (Q50, Q53 and Q109) neurons. Data represent mean ± SD; ****p<0.0001. **(J).** MEs isolated from control neurons (wtHTT-Q18, Q20 and Q33) and HD neurons (mHTT-Q50, Q53 and Q109) and analyzed by WBs to measure the PNKP protein levels in steady-state. TFAM was used as loading control. Lane M: protein molecular weight markers in kDa. **(K).** DNA isolated from control (Q18, Q20 and Q33; lanes 1 to 3) and HD iPSC-derived neurons (Q50, Q53 and Q109; lanes 4 to 6) and 8.9 kb and 0.2 kb segments of mtDNA PCR amplified, and the PCR products analyzed by agarose gel. Relative amplification efficiencies of the 8.9 kb and 0.2 kb segments of mtDNA shown by arrowheads. **(L).** Relative PCR amplification efficiencies of human mtDNA segments (8.9 kb and 0.2 kb) from the control vs. HD neurons. Data represents mean ± SD; ****p<0.0001.

To further validate our findings, we also measured mitochondrial PNKP activity and mtDNA repair efficacy in a well-characterized neuronal PC12 cell model of HD that recapitulates many aspects of cellular phenotypes of the disease[8]. We isolated pure mitochondria from wildtype neuronal PC12 cells and from PC12 cells expressing exogenous wtHTT-Q23 or mHTT-Q148[8]. We then assessed PNKP activity in the mitochondrial extracts by measuring its 3’-phosphatase activities. As expected, cells expressing exogenous HTT (HTT-Q23) showed ∼60-70% increased PNKP activity compared to control wildtype PC12 cells. In contrast, PNKP activity in the mitochondrial extracts from cells expressing mHTT (mHTT-Q148) was reduced by ∼60-70% compared with control PC12 cells (Supplementary figures 5A and 5B). Importantly, the steady state levels of PNKP protein in the MEs of mutant versus control cells remained unchanged (Supplementary figure 5C), indicating that HTT and mHTT enhance and decrease mitochondrial PNKP activity, respectively, but do not affect protein levels. These data further validate our findings that HTT is necessary for mtDNA repair efficacy, while mHTT impairs mtDNA repair, similar to its effects on nuclear DNA repair [21]. We next performed LA-QPCR analysis in mutant and control cells to confirm that reduced DNA repair activity results in compromised mtDNA integrity. Damage accumulation was ∼2 fold increased in mutant PC12 cells expressing mHTT-Q148 compared to controls (Supplementary figures 5D; lanes 3-4 vs. lanes 1-2, and 5E). Finally, increasing PNKP expression in mutant cells expressing mHTT-Q148 significantly rescued mtDNA repair efficacy (Supplementary figures 5D; lanes 5-6 vs. lanes 3-4, and 5E), reduced cell toxicity (Supplementary figure 5F), and improved mitochondrial membrane potential (Supplementary figure 5G). Overall, these experiments demonstrate a critical role for PNKP activity in maintaining mitochondrial genome integrity.

We next tested whether mHTT disrupts the interactions of PNKP with the macromolecular complex assembled by HTT by using SH-SY5Y cells overexpressing either HTT or mHTT (Supplementary figure 6A). Compared to control cells, HTT-overexpressing (HTT-OE) SH-SY5Y cells showed significantly increased interactions between HTT and POLRMT or PNKP as revealed by PLA analysis, and these PLA puncta overlapped with Mito-Tracker staining (Supplementary figures 6B to 6D), suggesting that increasing HTT levels results in increased assembly of the mtDNA repair complex. Furthermore, SH-SY5Y cells expressing exogenous HTT also showed significantly increased mtDNA repair activity, while cells expressing mHTT-Q145 showed ∼70% lower mtDNA repair activity compared with controls (Supplementary figures 6E and 6F). Consistent with our interpretation, cells expressing exogenous HTT-Q19 showed improved mtDNA integrity (Supplementary figures 6G; lanes 4-6 vs. lanes 1-3 and 6H) over mHTT-expressing cells which showed compromised mtDNA integrity (Supplementary figures 6G; lanes 7-9 vs. lanes 1-3, and 6H). Notably, increasing HTT levels in mutant cells dramatically reduced mtDNA damage and improved overall mtDNA integrity (Supplementary figures 6I; lanes 7-9 vs. lanes 4-6, and 6J).

To test the relevance of our studies to human disease, we investigated whether HD patient-derived cells have lower mtDNA repair activity by measuring PNKP activities in mitochondrial protein extracts from HD and unaffected control induced pluripotent stem cells (iPSCs)-derived striatal neurons [21, 38]. We found ∼70-80% reduction in PNKP activity in mitochondrial extracts from HD iPSC-derived striatal neurons (mHTT allele encoding Q50, Q53, or Q109) compared with controls (with Q18, Q20, and Q33 alleles) (Figures 5H and 5I). Additionally, consistent with our cell culture data (Figures 5A to 5C), WBs of the MEs from iPSC-derived neurons revealed no change in the steady-state levels of mitochondrial PNKP protein between the mutant and control cells (Figure 5J). The LA-QPCR analyses also revealed significantly increased mtDNA damage in HD neurons (Q50, Q53, and Q109) compared to control neurons (Q18, Q20, and Q33) (Figures 5K and 5L). Thus, analyses from multiple HD cell models, and from iPSC-derived striatal neurons from HD patients, strongly suggest that mHTT disrupts the activity of the mtDNA repair complex in neurons, which leads to increased mtDNA damage accumulation, and compromised mitochondrial function.

### Mutant HTT with extended polyglutamine sequences impairs mtDNA repair in mouse and *Drosophila* models of HD

Many studies suggest that DNA damage may be an early event in the progression of neurodegenerative diseases[39–41]. Therefore, we hypothesized that mtDNA damage occurs early during the progression of HD partly due to perturbation of the mtDNA repair mechanisms by mHTT. To test this hypothesis *in vivo*, we measured 3’-phosphatase activity of PNKP in MEs isolated from various brain regions, including the striatum (STR), cerebral cortex (CTX) and cerebellum (CRBL) of pre-symptomatic 7-week-old zQ175 transgenic mice expressing full-length mHTT[42] and age-matched wildtype controls. Neuropathological analyses has revealed that despite ubiquitous expression of mHTT in various brain regions, it impacts the STR and CTX more severely while relatively spares the CRBL[43]. Therefore, we tested whether mHTT expression impacts these brain regions differentially. Similar to the data from our cell culture models of HD, we found a ∼60-70% reduction of PNKP activity in the STR and CTX of zQ175 mice, and only ∼3-4% reduction of PNKP activities in the CRBL, with no change in steady-state PNKP protein levels (Figures 6A to 6C). Accordingly, LA-QPCR analysis showed greater mtDNA damage in the CTX and STR, but substantially less damage in the CRBL of zQ175 mouse brains at the same point (Figures 6D and 6E). These data suggest that mtDNA damage accumulation is an early event in the progression of HD, likely resulting from defective repair mechanisms that mostly affect the STR and CTX. Finally, to validate our findings in human disease, we isolated genomic DNA from post-mortem caudate nucleus tissue from HD patients and age-matched control subjects to measure relative levels of mtDNA damage. LA-QPCR analyses revealed the presence of substantially higher mtDNA damage in HD patients (n=7) compared with brain tissue from control subjects (n=3) (Figures 6H and 6I).

**Figure 6:**
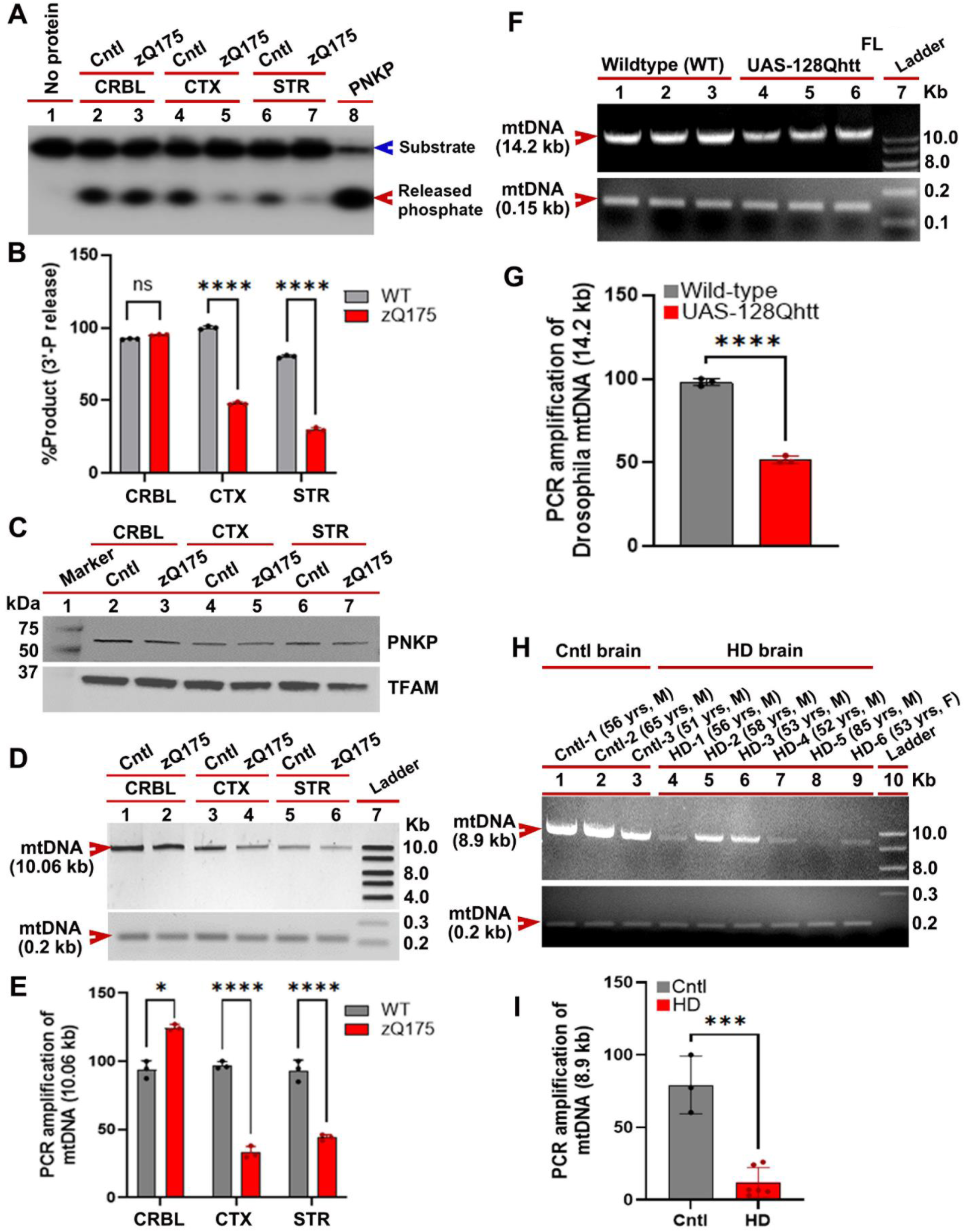
MtDNA damage accumulates in HD transgenic mouse, in transgenic *Drosophila* model of HD, and in HD patients’ brain. **(A).** MEs purified from the cerebellum (CRBL; lanes 2 vs. 3), cortex (CTX; lanes 4 vs. 5), and striatum (STR; lanes 6 vs. 7) of asymptomatic (7-weeks-old) heterozygous zQ175 male mice and age-matched control (Cntl) mice, and MEs added to the radio-labelled DNA substrate to measure the release of 3’-phosphate (red arrowhead) from the 32-P-labelled DNA substrate (blue arrowhead); Lane 1: No protein extract added to the DNA substrate as a negative control, and Lane 8: Purified PNKP protein (25 femtomoles) added to the substrate as a positive control. **(B).** Relative mitochondrial PNKP activities (in terms of % product) in CRBL (lanes 2 vs. 3 in figure 6A), CTX (lanes 4 vs. 5 in figure 6A), and STR (lanes 6 vs. 7 in figure 6A) of zQ175 transgenic male mice and age-matched control (Cntl) mice. Three biological replicates and three technical replicates used in this study. Data represents mean ± SD; ****p<0.0001; ns = not significant. **(C).** Mitochondria purified and the MEs isolated from the CRBL, CTX and STR from 7-week-old zQ175 transgenic and age-matched control (Cntl) mice and analyzed by WBs to determine PNKP protein levels. TFAM was used as loading control. Lane 1: protein molecular weight marker in kDa. **(D).** DNA isolated from the CRBL, CTX, and STR of asymptomatic (7-week-old) zQ175 transgenic and age-matched control (Cntl) mice. The 10.06 kb and 0.2 kb sections of mtDNA PCR amplified, and the PCR products from control (lanes 1, 3, and 5) and zQ175 transgenic mice (lanes 2, 4, and 6) analyzed on agarose gels. The 10.06 kb and 0.2 kb segments of mtDNA PCR-amplified and PCR products are analyzed on agarose gels and by red arrowheads. Lane 7: 1-kb DNA ladder. **(E).** Relative PCR amplification efficacies of mtDNA segments in age-matched controls (Cntl) and 7-week-old heterozygous zQ175 mouse brains. Three biological replicates and three technical replicates used in this study. Data represent mean ± SD; *p<0.05 and ****p<0.0001. **(F).** DNA isolated from the UAS-128Qhtt^FL^ transgenic *Drosophila* and age-matched wildtype (WT) control fly heads. The 14.2 kb and 0.15 kb sections of *Drosophila* mtDNA PCR amplified, and the products from wildtype (WT) control (lanes 1 to 3) and UAS-128Qhtt^FL^ transgenic flies (lanes 4 to 6) analyzed on agarose gels. *Drosophila* mtDNA segments shown by red arrowheads. Twenty-five biological replicates and three technical replicates were used for this measurement. Lane 7: 1-kb DNA ladder. **(G).** Relative PCR amplification efficacies of *Drosophila* mtDNA segments in wildtype control and 20 days-old UAS-128Qhtt^FL^ *Drosophila* brains. Twenty-five biological replicates and three technical replicates were used in this study. Data represent mean ± SD; ****p<0.0001. **(H).** Total DNA isolated from the caudate of three controls (Cntl; lanes 1-3) and 6 HD patients’ brains (lane 4: 56 years-old, male; grade 2; lane 5: 58 years-old, male; grade 2; lane 6: 53 years-old, male; grade 3; lane 7: 52 years-old, male; grade 3; lane 8: 85 years-old, male; grade 3; lane 9: 53 years-old, female; grade 3). The 8.9 and 0.2 kb sections of mtDNA PCR amplified, and the PCR products from the control subjects (lanes 1 to 3) and HD patients (lanes 4 to 9) analyzed on agarose gels (arrowheads). Five biological replicates and three technical replicates used in this study. Lane 10: 1-kb DNA ladder. **(I).** Relative PCR amplification efficacies of various mtDNA segments from brain tissue from HD patients (HD) and age-matched control subjects (Cntl). Five biological replicates and three technical replicates used in this analysis. Data represent mean ± SD; ***p<0.001.

*Drosophila* has long been used to model various neurological diseases, including HD[44–47]. Transgenic flies expressing human full-length mHTT in neurons recapitulate many HD-like phenotypes, including reduced longevity, motor deficits, and progressive neurodegeneration[48]. Since mitochondrial abnormalities have also been observed in *Drosophila* models of HD [49], we assessed mtDNA integrity in a *Drosophila* HD model. Specifically, we tested whether neuronal expression of mHTT causes mtDNA damage in neurons by assessing mtDNA integrity in *Drosophila* brains expressing full-length human mHTT (UAS-128Qhtt^FL^)[49]. Consistent with our data from cell culture and mouse models, LA-QPCR analyses showed significantly higher mtDNA damage in UAS-128Qhtt^FL^ transgenic fly brains compared with age-matched control (UAS-16Qhtt^FL^) flies (Figures 6F and 6G).

While our data suggests that full-length mHTT is present within mitochondria, previous studies have demonstrated that N-terminal truncated fragments of mHTT are present in HD brain, resulting from proteolytic cleavage and inappropriate splicing[50, 51]. These truncated N-terminal fragments of mHTT (NT-mHTT) are highly toxic and cause neurodegeneration[52], and NT-mHTT has been found within mitochondria[53]. Therefore, we tested whether mitochondrial NT-mHTT fragment interferes with mitochondrial PNKP activity and mtDNA repair in two well-characterized HD transgenic mouse models that express the N-terminal region of mHTT: N171-82Q[53] and R6/2[53] mouse lines. First, we measured mitochondrial PNKP activity in MEs isolated from CTX, STR and CRBL of the N171-82Q transgenic mouse brains expressing NT-mHTT containing 82 glutamine repeats[54].We observed >70% reduction in PNKP activity in MEs from the CTX and STR of asymptomatic (7-week-old) heterozygous N171-82Q mice, with only marginally lower PNKP activity in the CRBL (Supplementary figures 7A and 7B). Consistently, the N171-82Q CTX and STR, but not the CRBL, showed substantially more mtDNA damage than age-matched control mice as revealed by LA-QPCR analyses (Supplementary figures 7C and 7D). We next measured the mitochondrial PNKP activity and mtDNA damage in 3-month-old R6/2 HD transgenic mouse brain expressing NT-mHTT[55]. At these early time-points, PNKP activity in MEs from the CTX and STR of heterozygous R6/2 mice was substantially decreased (by >80%), with no significant decrease in the CRBL, compared with controls (Supplementary figures 7E and 7F). The R6/2 CTX and STR also showed substantially more mtDNA damage than controls, while mtDNA damage was minimal in the CRBL (Supplementary figures 7G and 7H). Thus, results from various HD models and post-mortem HD brains support the hypothesis that expression of either full-length mHTT or the N-terminal truncated fragment of mHTT causes a substantial decrease in mtDNA repair efficacy, likely occurring early in the disease progression.

### Restoring mtDNA repair in a *Drosophila* HD model improves mitochondrial genome integrity and rescues motor defects

Our data demonstrates that increasing HTT levels in cell culture models of HD restores mtDNA integrity and mitochondrial function that are compromised by mHTT. We therefore investigated whether increasing the stochiometric ratio of HTT to mHTT might increase the number of functional DNA repair complexes assembled by HTT within mitochondria and enhance overall mtDNA repair activity. Indeed, additional expression of exogenous full-length human HTT in mutant STHdhQ111 striatal cells dramatically improved mitochondrial PNKP activity (Figures 7A and 7B), and enhanced mtDNA repair and reduced mtDNA damage (Figures 7C and 7D). Since STHdhQ111 cells exhibit impaired mitochondrial function[56], we tested whether increasing HTT levels in STHdhQ111 cells would also improve the mitochondrial function deficits caused by mHTT[18, 57]. We treated control STHdhQ7, STHdhQ111, and STHdhQ111 cells expressing exogenous full-length HTT with the complex II inhibitor 3-NP in the presence of JC-10 dye. In polarized cells, JC-10 predominantly concentrates in the mitochondrial matrix, where it forms red/orange aggregates. In contrast, in depolarized cells, JC-10 diffuses out of mitochondria and exists in monomer that stain cells green. The mutant STHdhQ111 cells showed lower orange fluorescence compared to control STHdhQ7 cells, which is indicative of reduced mitochondrial membrane potential. Importantly, enhancing HTT levels in mutant STHdhQ111 cells significantly increased orange fluorescence, indicating restoration of mitochondrial membrane potential (Figure 7E).

**Figure 7.**
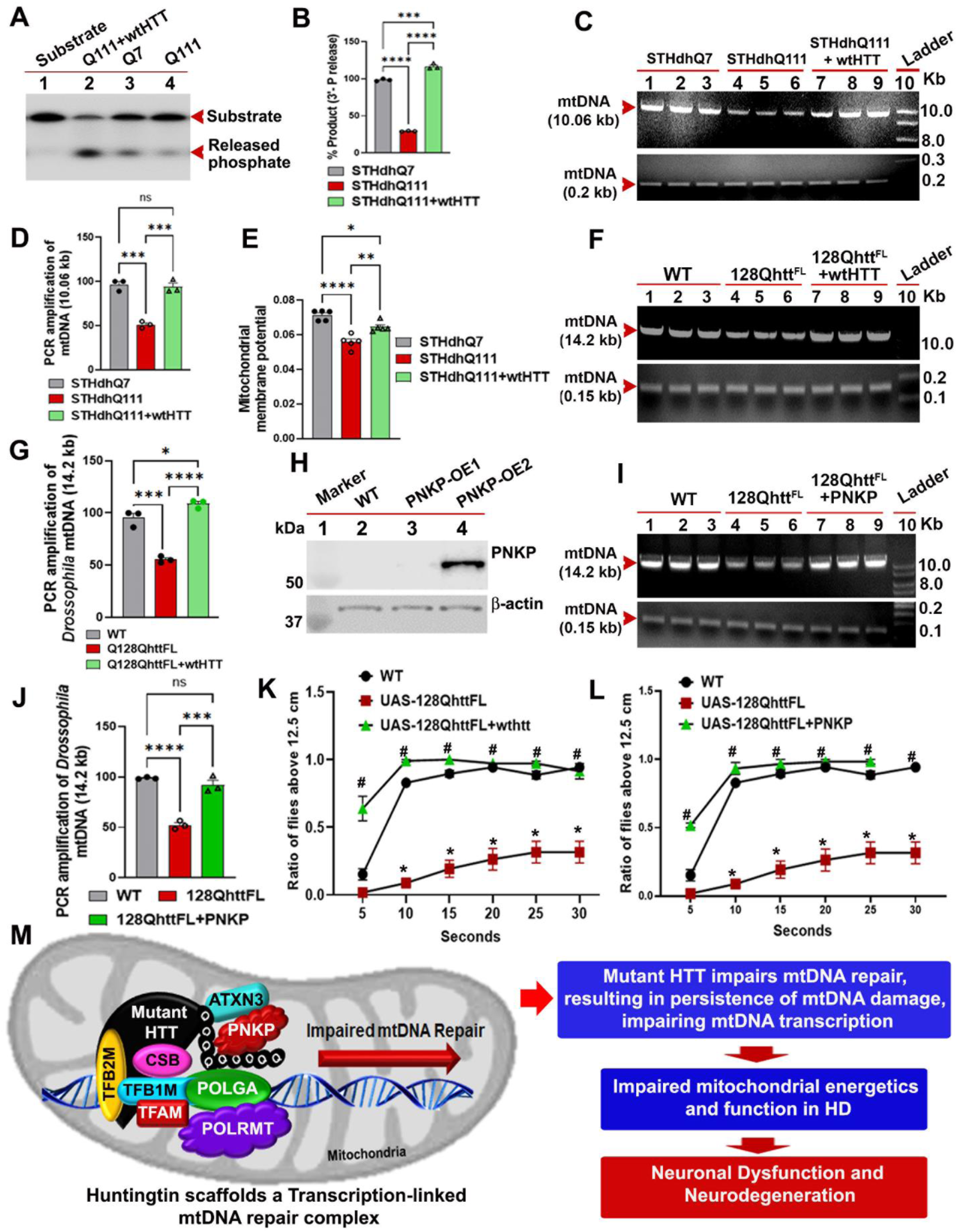
Restoring mtDNA repair activity in a *Drosophila* model of HD improves mtDNA integrity, and mitochondrial function and motor defects. **(A).** PNKP activities in MEs are isolated from the control STHdhQ7 cells (lane 3; Q7), mutant STHdhQ111 cells (Q111; lane 4) and STHdhQ111 cells overexpressing exogenous wtHTT (Q111 + wtHTT; lane 2); Lane 1: no protein was added to the DNA substrate as negative control. The DNA substrate and released phosphate bands shown with arrowheads. **(B).** Relative mitochondrial PNKP activities (in terms of % product) in STHdhQ7, STHdhQ111 and STHdhQ111 cells overexpressing wtHTT. Three biological and three technical replicates were used in this study. Data represent mean ± SD; ****p<0.0001; ***p<0.001. **(C).** Total genomic DNA isolated from STHdhQ7 (lanes 1 to 3), STHdhQ111 (lanes 4 to 6) and STHdhQ111 cells overexpressing wtHTT (lanes 7 to 9), and 10.6 kb and 0.2-kb segments of mouse mtDNA PCR amplified; and the PCR products electrophoresed on agarose gels and stained with EtBr (red arrowheads). Lane 10: 1-kb DNA ladder. **(D).** Relative PCR amplification of mtDNA in STHdhQ7, STHdhQ111, and STHdhQ111 cells overexpressing wtHTT. Three technical replicates were used for the calculation. Data represents mean ± SD, ***p<0.001; ns = not significant. **(E).** Control wildtype STHdhQ7 cells, mutant STHdhQ111 cells, and mutant STHdhQ111 cells expressing exogenous wtHTT treated with JC-10 dye, and mitochondrial membrane potential measured. Five technical replicates were used for calculations. Data represent mean ± SD; *p<0.05. **p<0.01; ****p<0.0001. **(F).** Total genomic DNA isolated from wildtype fly heads (WT; lanes 1 to 3), HD transgenic UAS-128Qhtt^FL^ fly heads (lanes 4 to 6) and the UAS-128Qhtt^FL^ fly heads expressing human wtHTT (lanes 7 to 9), and 14.2 kb and 0.15-kb segments of *Drosophila* mtDNA PCR amplified; and the products analyzed on agarose gels (red arrowheads). Lane 10: 1-kb DNA ladder. **(G).** Relative PCR amplification of mtDNA in control wildtype flies (WT), mutant UAS-128Qhtt^FL^, and mutant UAS-128Qhtt^FL^ *Drosophila* flies ectopically expressing human wtHTT. Twenty-five biological replicates and three technical replicates were used for this measurement. Data represent mean ± SD; *p<0.05; ***p<0.001; ****p<0.0001. **(H).** Total protein isolated from wildtype (WT) *Drosophila* heads (n=30; lane 2), and transgenic *Drosophila* heads expressing human PNKP, and effective expression of PNKP assessed by WBs. Transgenic line (PNKP-OE2; lane 4) showing effective transgenic expression of human PNKP in heads. Lane 3: Transgenic line PNKP-OE1 does not show effective expression of human PNKP. Lane 1: Protein molecular weight marker in kDa. **(I).** Total DNA isolated from wildtype fly heads (WT; lanes 1 to 3), HD transgenic UAS-128Qhtt^FL^ fly heads (lanes 4 to 6) and the UAS-128Qhtt^FL^ fly heads expressing human PNKP (lanes 7 to 9), and 14.2 kb and 0.15-kb sections of *Drosophila* mtDNA PCR amplified; and the PCR products analyzed on agarose gels (red arrowheads). Lane 10: 1-kb DNA ladder. **(J).** Relative PCR amplification of mtDNA in control wildtype flies (WT), mutant UAS-128Qhtt^FL^, and mutant UAS-128Qhtt^FL^ flies ectopically expressing human PNKP. Twenty-five biological replicates and three chemical replicates were used for this measurement. Data represent mean ± SD; ***p<0.001; ****p<0.0001; ns = not significant. **(K).** Climbing properties of 20-day-old *Drosophila* that reflect the motor coordination in wildtype, UAS-128Qhtt^FL^ and UAS-128Qhtt^FL^ flies expressing exogenous wtHTT measured by standard climbing properties of flies. Relative motor function of wildtype (WT), UAS-128Qhtt^FL^, and UAS-128Qhtt^FL^ Drosophila expressing exogenous wtHTT are shown. *p<0.001 (UAS-128Qhtt^FL^ vs. WT) and <00.1 (mutant flies expressing exogenous wtHTT vs. mutant flies). **(L).** Relative motor function of wildtype control (WT), HD transgenic (UAS-128Qhtt^FL^) and in HD transgenic flies ectopically expressing human PNKP. *p<0.001 (UAS-128Qhtt^FL^ vs. WT) and # p<00.1 (mutant flies expressing exogenous PNKP vs. mutant flies). **(M).** Schematic diagram of our hypothesized mechanism by which mHTT with polyQ expansions compromises mtDNA repair and integrity, and function in HD. Wildtype native HTT forms a mtDNA repair complex with POLRMT, TFAM, DNA POLGA, CSB, TFB1M/TFB2M, and PNKP, and this multi-protein complex edits mtDNA lesions during transcription to preserve mtDNA integrity, replication, and transcription. In HD, mHTT impairs the normal function of this complex, leading to persistence of mtDNA damage. This putative pathway explains how mHTT adversely impacts mtDNA repair, and perturbs mtDNA gene transcription to disrupt mitochondrial homeostasis and function to trigger early neurotoxicity in HD.

Finally, to determine whether increasing wtHTT expression in an *in vivo* HD model would improve mtDNA repair, and mitochondrial genome integrity, we again turned to the *Drosophila* model of HD. Human HTT (16Qhtt^FL^) and mHTT (128Qhtt^FL^) were expressed in *Drosophila* neurons using the UAS-Gal4 system[58]. We used the BG-380 gal4 driver[59] to specifically drive expression of UAS-16Qhtt^FL^ in the neurons of the 128Qhtt^FL^ flies. The fly heads were separated on dry ice, total DNA was isolated, and mitochondrial genome integrity was assessed by performing LA-QPCR. Consistent with our cell culture data, expression of mHTT led to an increase in mtDNA damage compared to wildtype flies. Importantly, increasing HTT levels in mHTT background (UAS-128Qhtt^FL^) substantially restored mitochondrial genome integrity (Figures 7F and 7G), indicating that HTT enhances mtDNA repair *in vivo*. However, since HTT is known to function in many cellular processes, including endocytosis, ciliogenesis, intracellular transport of organelles and vesicles in the axons[60–62], we tested whether stimulating mtDNA repair was sufficient to restore mtDNA integrity *in vivo*. We therefore generated a transgenic fly line expressing human PNKP, which robustly expressed human PNKP when driven by Gal4 (Figure 7H). To evaluate whether increased expression of PNKP could rescue the mtDNA repair deficiency caused by mHTT, we co-expressed the PNKP transgene with mHTT in fly neurons using the neuron-specific BG-380 Gal4 driver[59]. Remarkably, PNKP expression significantly restored mtDNA integrity in these flies, indicating that the mtDNA repair deficiency was due to decreased PNKP activity, consistent with our cell culture findings (Figures 7I and 7J).

Next, we examined whether restoring the mtDNA repair could improve the motor deficits commonly associated with fly models expressing mHTT[48] by raising flies expressing either HTT, mHTT, flies co-expressing HTT and mHTT and flies co-expressing PNKP and mHTT. Motor function was assessed using the standard negative geotaxis climbing assay[63], which has previously demonstrated that expression of mHTT in fly neurons causes motor deficits after ∼ 20 days post-eclosion[48]. Consistent with these studies, flies expressing mHTT exhibited severe motor deficits after 20 days. However, flies co-expressing mHTT with either HTT or PNKP showed a marked improvement in motor function, with most flies able to climb comparably to flies expressing wtHTT alone (Figures 7K and 7L). These results strongly suggest that enhancing mtDNA repair efficacy *in vivo* improves both mtDNA integrity and motor phenotypes associated with HD. Together, these data indicate that mHTT compromises the efficacy of mtDNA repair, likely through direct interference with mtDNA repair mechanism(s). Further, increasing either the ratio of HTT to mHTT, or restoring PNKP activity, can rescue the defects in mtDNA repair and neuronal function in both cell culture and *in vivo* models. In summary, our data provides a mechanistic explanation for the neuroprotective effects of HTT expression in mitochondria[2–4, 6], and why mHTT induces mtDNA damage and causes extensive mitochondrial dysfunction and degeneration in HD[10, 14, 15, 18, 24, 64].

## DISCUSSION

Over the past three decades, research has established that mitochondrial dysfunction and metabolic deficiencies precede the onset of neurological symptoms and overt neurodegeneration in HD[65], highlighting their critical role in disease initiation and progression. Remarkably, mitochondrial dysfunction in HD was recognized even before the discovery of the HTT gene[66]. Impaired mitochondrial function appears early and persists chronically, as evidenced by progressive weight loss despite normal caloric intake[51, 67], abnormal mitochondrial morphology, and decreased electron transport chain activity in postmortem HD brain tissues[16, 66, 68]. Mouse models of HD also consistently demonstrate impaired mitochondrial respiration[69–72]. Furthermore, accumulation of oxidized bases and mtDNA deletions are frequently observed in both animal models and postmortem human brain tissue[69–72]. These findings strongly implicate defective mtDNA repair and subsequent accumulation of mtDNA lesions as contributors to mitochondrial dysfunction in HD.

Mitochondria respond to nuclear DNA damage repair signaling through NADH sensing, and oxidative stress from mitochondrial dysfunction or inflammatory responses compromises DNA integrity and repair. Several nuclear repair enzymes and adaptors—such as EXO5 and XRCC1—also localize to mitochondria[73–77]. A growing body of evidence suggests that wtHTT plays an essential role in maintaining mitochondrial function. For instance, reducing wtHTT levels in embryonic stem cells disrupts mitochondrial structure, distribution, and function[8, 78]. Knockdown of wtHTT in the postnatal brain results in rapidly progressive neurodegeneration[7], while its overexpression provides significant neuroprotection in vivo and in vitro[2–4, 6]. However, the mechanisms through which wtHTT supports mitochondrial function, and how mHTT disrupts it, remain poorly defined.

Our study employs immunoprecipitation under stringent conditions, imaging, ChIP, and ChIP-re-ChIP analyses to demonstrate that wtHTT localizes within mitochondria, forms a multiprotein complex with POLRMT, POLGA, DNA repair enzymes, and mitochondria-specific transcription factors, and interacts directly with mtDNA. We propose that this novel complex facilitates transcription-coupled mtDNA repair, analogous to the TCR complex in the nucleus [21]. Prior studies identified TCR-initiator proteins, including Cockayne syndrome (CS) proteins A and B (CSA and CSB), in mitochondria[28, 29]. These proteins are increasingly recruited to mitochondria upon exposure to DNA-damaging agents[29]. Given the essential role of CS proteins in initiating nuclear TCR[32–34] and their involvement in mtDNA repair[28, 29], our discovery of CSB, POLRMT, and POLGA in the HTT-organized complex supports our hypothesis that mitochondria utilize a transcription-coupled repair mechanism akin to nuclear TCR.

We show that elevated wtHTT expression significantly enhances mtDNA repair efficacy, partly by boosting PNKP activity, a crucial enzyme for both nuclear and mtDNA repair[21, 27, 79–81]. This enhanced repair capacity may explain the neuroprotective effects of wtHTT[82]. In contrast, mHTT binds to the same protein complex as wtHTT—potentially with greater affinity— disrupting mtDNA repair and contributing to the accumulation of lesions, a hallmark of mitochondrial dysfunction in HD[15–17].

### Protective role of HTT in Mitochondria

HTT depletion in murine stem cells leads to mitochondrial degeneration and dysfunction[8]. Subcellular fractionation[11] and imaging[83] confirm a strong association between HTT and mitochondria. Multiple studies report HTT localization to brain mitochondria[12, 84]. However, many investigations into mHTT’s effects on mitochondrial function relies on *in vitro* assays or exogenously expressed HTT-fusion proteins[12, 84], making it challenging to conclusively determine wtHTT’s presence in mitochondria. Although evidence suggests that wtHTT maintains mitochondrial function and mHTT impairs it[14, 15, 18, 24], the precise mitochondrial role of HTT remains unclear.

Native wtHTT associates with brain mitochondria in both wild-type and HD transgenic mice[12, 85, 86]. The first 17 amino acids of wtHTT are required for localization of exon1 HTT to mitochondria[83]. mHTT may associate with the outer mitochondrial membrane, disrupt protein transport across membranes, and interact with DRP-1, a protein critical for mitochondrial quality control[11–13]. While our study highlights the importance of mtDNA repair in supporting mitochondrial function, additional research is needed to clarify how wtHTT maintains mitochondrial integrity and how it is imported into mitochondria. The first 17 amino acids of wtHTT may facilitate mitochondrial localization[83], but this requires rigorous validation. Further sub-mitochondrial localization studies are also necessary to explore potential additional functions of wtHTT[15–17].

### HTT Organizes a Macromolecular mtDNA Repair Complex

Our findings position wtHTT as a key organizer of a DNA repair and transcription complex within mitochondria. Given wtHTT’s role in nuclear transcription-coupled repair[21], we hypothesize that it may also scaffold a novel mitochondrial TCR complex[87]. This is supported by evidence that wtHTT binds core components of the DNA repair machinery, mitochondrial RNA polymerase (POLRMT), mitochondrial DNA polymerase, and transcription factors. We suggest that mHTT’s binding to this complex disrupts its function, causing it to stall at damaged sites in the mutant cells. This impairment of mtDNA repair and transcription may abort mitochondrial gene expression or trigger signals for mtDNA degradation and mitophagy[88].

Future research should examine whether mitochondrial gene transcription is disrupted in HD and at what stage of disease progression.

### Conclusions

Overall, our data supports a novel mitochondrial HTT function i.e., to organize a transcription-linked DNA repair complex that maintains mitochondrial genome integrity and thereby mitochondrial function. The results offer key insights into how increasing wtHTT expression in neurons may provide neuroprotection[2–4, 6]; why depletion of wtHTT in mouse brain triggers neurotoxicity[7]; and how mHTT may cause mitochondrial dysfunction in the early phases of HD[10, 14, 15, 18, 24, 64]. Collective findings highlight the importance of wtHTT in maintaining mtDNA integrity and normal mitochondrial function in neurons. Therefore, genetic therapies with anti-sense oligos (ASOs) directed toward lowering the mutant copy of HTT in HD patients, which also reduced the wtHTT[89], led to devastating consequences and manifested with motor and memory deficits[90]. However, it remains to test whether selectively and allele-specifically lowering of mHTT *in vivo* reduces mtDNA damages and improves the motor and memory deficits in HD patients. Based on our current findings, we reason that either allele-specific knockdown of the mHTT or increasing wtHTT may offer an alternative and meaningful strategy for improving DNA repair efficacy and for rescuing the motor and memory defects in HD.

### Limitation of this study

Although data presented here show how wtHTT may support efficient mtDNA repair, it does not explain selective vulnerability of cells and brain regions in HD. The mtDNA repair complexes could consist of different proteins in different brain regions, and inactivation or activation of such “region specific” factors might be differentially regulated, giving rise to differential susceptibility. These putative factors could be either protein modifying enzymes (post-translational modification), e.g., Protein Inhibitor of Activated STAT (PIAS1) which can SUMO modify PNKP[91], or RNA processing factors that might synchronously regulate transcription, DNA repair and RNA processing. It may prove useful to investigate whether additional factors are present within the putative mitochondrial TCR complex that could regulate its activity. Alternatively, certain brain regions might be differentially vulnerable to the effects of mtDNA damage due to differential energy requirements. Thus, studies that focus on the elucidation of the composition of the mtDNA damage repair complex, on the putative mitochondrial TCR complex in specific brain regions, and on understanding how its activity is regulated, are expected to provide further insight into the selective vulnerability in HD.

## ACKNOWLEDGEMENTS

This research was supported by National Institutes of Health grants RO1 NS130830 and to R01 EY026089-01A1 to P.S.S., and Hereditary Disease Foundation (HDF) Grant to P.S.S; R01-NS100529 to L.M.E.; PO1 CA092584 and R35 CA 220430 to J.A.T; R01 AG033082 and R01 NS065874 to A.R.L.S., R35 NS116872 and R01 NS090390 to L.M.T; a Hereditary Disease Foundation Fellowship to C.G; a Robert Welch Chemistry Chair G-0010 to J.A.T.; R56 NS105681, Alzheimer’s Association grant to Y.P.W., Alzheimer’s Association grant AARF-22-967275 to S.G. and the Nancy and Buster Alvord Endowment to C.D.K.

## AUTHOR CONTRIBUTIONS

S.P., S.G., N.Z., T.K.P. generated ChIP, ChIP-re-ChIP, WB, IP, PLA, and LA-QPCR analyses. S.P. and P.S.S. generated various plasmid constructs and cell models of HD, performed LA-QPCR, and quantitative PCR analyses. N.Z. performed the cell toxicity assays, and mitochondrial membrane potential measurements and helped draft the manuscript. C.S-G., and L.M.T. characterized and provided the iPSC-derived primary neurons, provided brain tissues from HD mouse models, and helped draft the manuscript. E.L.M. helped draft the manuscript. K.B and Y.P.W. generated the *Drosophila* expressing human PNKP and performed the motor function analysis and helped draft the manuscript. S.Y. performed the confocal image analyses and A.C. and T.K.H. performed the PNKP assays. A.S.D., and A.R.L.S. characterized the HD transgenic mouse lines, processed the brain tissue, provided the tissues for various analyses, and helped draft the manuscript. T.A., and C.D.K. provided data interpretation and helped draft the manuscript. L.M.E. generated and provided the PC12 inducible cells, zQ175 transgenic mouse tissue and various cDNA clones expressing wildtype and mutant HTT. C-L T, and J.A.T. did computational modeling plus contributed to analyzing interfaces and writing the manuscript. P.S.S. was the project leader and wrote and edited the final manuscript with substantial input from L.M.T., J.A.T.

## SUPPLEMENTARY FIGURE LEGENDS

**Supplementary figure 1: HTT forms macromolecular complex with the mitochondrial transcription complex components in mitochondria**.

**(A).** PLA in control (-GO) SH-SY5Y cells with anti-HTT mouse monoclonal Ab (MAB2170; Millipore-Sigma) and anti-POLRMT rabbit polyclonal Ab (GTX105137; Genetex).

**(B).** PLA in GO treated SH-SY5Y cells with an anti-HTT mouse monoclonal Ab (MAB2170; Millipore-Sigma) and with anti-POLRMT rabbit polyclonal Ab (GTX105137; Genetex).

**(C).** Relative PLA signals showing substantial increased interaction of endogenous HTT with POLRMT within the mitochondria in GO-treated (+GO) cells compared to control (-GO). Data represent mean ± SD; ****p<0.0001.

**(D).** Plasmids encoding FLAG-tagged wtHTT-Q24 and Myc-tagged PNKP, POLRMT, POLGA or TFAM separately co-transfected into human HEK293 cells, cell-extracts isolated 48 hours post-transfection, and analyzed by WBs to detect the expressions of exogenous FLAG-tagged HTT and Myc-tagged PNKP, -POLRMT, -POLGA, and -TFAM (arrows).

**(E).** Cell extracts isolated from the co-transfected HEK293 cells expressing FLAG-tagged HTT-Q24 (FLAG-HTT-Q24), and Myc-tagged PNKP, POLRMT, POLGA or TFAM. The extracts were IP’d with anti-FLAG Ab (F3165; Sigma), and FLAG ICs analyzed by WBs to detect exogenous Myc-tagged-POLRMT, -POLGA, -PNKP or - TFAM. IgG heavy chain and light chain shown by arrows. Lane 1: Protein molecular weight marker.

**(F).** MEs isolated from SH-SY5Y cells constitutively expressing exogenous (Ex) FLAG-tagged HTT-Q24, and IP’d with anti-FLAG Ab (F3165; Millipore-Sigma). The FLAG IC analyzed by western blotting to detect endogenous (En) POLRMT, POLGA, PNKP. Lane 1: protein molecular weight marker; lane 2: Input; lane 3 IgG IP; lane 4: FLAG IP and lane 5: total cell extract (Total CE).

**Supplementary figure 2: PNKP and CSB are present in mtDNA repair complex.**

Proximity ligation assay (PLA) performed to assess interaction/association of PNKP and mitochondrial transcription factor TFAM, or mitochondrial RNA polymerase (POLRMT) and Cockayne syndrome protein B (CSB) in mitochondria in SH-SY5Y cell to examine possible protein interactions. Mitochondria were stained with Mito Tracker Red (Panels 2), and green fluorescence indicates positive protein-protein interactions (arrows in panels 1 through 4 indicate representative positive PLA signals).

**A**). PLA with anti-TFAM rabbit monoclonal (ab176558; Abcam) and anti-PNKP mouse monoclonal Abs in SH-SY5Y cells without GO treatment.

**(B).** SH-SY5Y cells treated with GO (300 ng/mL) for 30 minutes for induction of mtDNA damage, and PLAs were performed with anti-TFAM rabbit (ab176558; Abcam) and anti-PNKP mouse Abs. Nuclei stained by DAPI. Positive signals indicated by arrows.

**(C).** Relative PLA signals showing interaction of TFAM and PNKP within mitochondria in control (-GO) cells and GO-treated (+ GO) cells. Data represent mean ± SD; ****p<0.0001.

**(D).** PLA with anti-POLRMT rabbit (GTX105137; Genetex) and anti-CSB mouse (sc-398022; Santa Cruz Biotech) Abs in control SH-SY5Y cells.

**(E).** SH-SY5Y cells treated with GO (300 ng/mL) for 30 minutes, and PLAs performed with anti-POLRMT rabbit (GTX105137; Genetex) and anti-CSB mouse (SC-398022; Santa Cruz Biotech) Abs.

**(F).** Relative PLA signals showing interaction of POLRMT and CSB within mitochondria in control (-GO) cells and GO-treated (+ GO) cells. Data represent mean ± SD; ****p<0.0001.

**(G).** Bimolecular fluorescent complementation (Bi-FC) experiment. Plasmids p-FLAG-VN-HTT and p-HA-VC-POLRMT, expressing full-length wtHTT (FLAG-tagged) and POLRMT (HA-tagged) respectively co-transfected in SH-SY5Y cells, and fluorescence reconstitution (arrowheads) assessed with confocal image analysis. Nuclei stained with DAPI.

**(H).** Plasmids p-FLAG-VN-HTT and p-HA-VC-POLGA, expressing the full-length wtHTT-Q19 (FLAG-tagged) and POLGA (HA-tagged), respectively co-transfected in SH-SY5Y cells, and fluorescence reconstitution (arrowheads) assessed with confocal image analysis. Nuclei stained with DAPI.

**Supplementary figure 3: HTT forms a mitochondria-specific DNA repair complex with mitochondrial POLRMT and POLGA in vivo.**

Proximity ligation assay (PLA) performed to assess possible interaction/association of HTT with mitochondrial RNA polymerase (POLRMT) or mtDNA polymerase gamma (POLGA) in human brain section (56 years; basal ganglia) to examine protein interactions. Mitochondria stained with anti-MT-Co1 antibody (panels 1), and green/yellow fluorescence indicates positive protein-protein interactions (arrows in panels 1 through 4 indicate representative positive PLA signals).

**(A).** PLA with anti-POLRMT (rabbit) and anti-HTT (goat) Abs.

**(B).** PLA with anti-POLGA (rabbit) and anti-HTT (goat) antibodies. Nuclei stained by DAPI.

**Supplementary figure 4. Disruption of mtDNA repair complex activity induces mtDNA damage in neuronal SH-SY5Y cells.**

**(A).** SH-SY5Y cells (lane 1), SH-SY5Y cells either expressing control-RNAi (lane 2) or POLRMT-RNAi (lane 3) differentiated, and MEs isolated and analyzed by WBs to detect POLRMT protein levels; Mitochondrial transcription factor TFAM used as loading control.

**(B).** Relative POLRMT protein levels (normalized to TFAM) in differentiated SH-SY5Y cells, SH-SY5Y cells expressing either control-RNAi or POLRMT-RNAi. Three biological and three technical replicates were used. Data represent mean ± SD; ****p<0.0001; ns = not significant.

**(C).** 3’-phosphatase activities of PNKP in the MEs from control SH-SY5Y cells (untreated; lane 1), cells expressing control-RNAi (Cntl-RNAi: lane 2), or POLRMT-RNAi (lane 3) determined by measuring the release of 3’ phosphate (red arrowhead) from the radio-labelled DNA substrate (blue arrowhead). Lane 4: Purified PNKP (25 femtomoles) added to the DNA substrate as a positive control; Lane 5: No protein extract added to the DNA substrate as negative control.

**(D).** Relative PNKP 3’-phosphatase activities (in terms of % product) in wildtype SH-SY5Y cells and SH-SY5Y cells expressing either the control-RNAi or POLRMT-RNAi. Data represent mean ± SD; ****p<0.0001; ns = not significant.

**(E).** DNA isolated from wildtype differentiated SH-SY5Y cells and SH-SY5Y cells expressing either control-RNAi (Cntl-RNAi) or POLRMT-RNAi (lane 3). 8.9 kb and 0.21 kb segments of mtDNA PCR amplified, and the PCR products analyzed on agarose gels. Lane 1: control SH-SY5Y cells (untreated); lane 2: cells expressing control-RNAi (Cntl-RNAi); lane 3: cells expressing POLRMT-RNAi; Lane 4: 1-kb DNA ladder.

**(F).** Relative mtDNA damages in differentiated control SH-SY5Y cells (untreated) and in SH-SY5Y cells expressing either control-RNAi (Cntl-RNAi) or POLRMT-RNAi. *p<0.05; **p<0.01; ns= not significant.

**(G).** SH-SY5Y cells (untreated; lane 1), and SH-SY5Y cells expressing either control-RNAi (Cntl-RNAi; lane 2) or PNKP-RNAi (lane 3) differentiated, and MEs isolated and analyzed by WBs to detect mitochondrial PNKP levels; TFAM used as loading control.

**(H).** Relative PNKP levels (normalized to TFAM) in SH-SY5Y cells (untreated) and SH-SY5Y cells expressing either control-RNAi or PNKP-RNAi. Data represent mean ± SD: ****p<0.0001; NS = not significant.

**(I).** DNA isolated from untreated SH-SY5Y cells, and SH-SY5Y cells expressing either PNKP-RNAi or control-RNAi (Cntl-RNAi). The 8.9 kb and 0.21 kb sections of mtDNA PCR amplified, and the PCR products analyzed on agarose gels and stained with EtBr (red arrowheads) and quantified. Lane 1: Untreated SH-SY5Y cells; lane 2: cells expressing control-RNAi; lane 3: expressing PNKP-RNAi, lane 4: 1-kb DNA ladder.

**(J).** Relative mtDNA damages in control SH-SY5Y cells (untreated), and cells expressing either control RNAi or PNKP-RNAi. Data represent mean ± SD; **P<0.01; ***p<0.001; ns = not significant.

**Supplementary figure 5. Increasing PNKP activity in PC12 cell model of HD rescues mtDNA integrity and mitochondrial physiology.**

**(A).** PNKP’s 3’-phosphatase activities in MEs (250 ng each) of control PC12 cells (Cntl: lane 1), PC12 cells expressing wtHTT-Q23 (lane 2), mHTT-Q148 (lane 3) or co-expressing mHTT-Q148 and exogenous PNKP (lane 4) determined by adding MEs to the radiolabeled DNA substrate and measuring the amount of radioactive phosphate release (red arrowhead) from the DNA substrate (blue arrowhead). Lane 5: Purified PNKP (25 femtomoles) added to the substrate as positive control, and Lane 6: no MEs added to the DNA substrate as negative control.

**(B).** Relative 3’-phosphatase activities of PNKP (in terms of % product) in MEs from wildtype PC12 cells (Cntl), PC12 cells expressing wtHTT-Q23, cells expressing mHTT-Q148 and co-expressing mHTT-Q148 and exogenous PNKP. Three biological and three technical replicates were used for all measurements. Data represent mean ± SD: ****p<0.0001; ns = not significant.

**(C).** MEs isolated from control PC12 cells (Cntl; lane 2), PC12 cells expressing wtHTT-Q23 (lane 3), mHTT-Q148 (lane 4) or co-expressing mHTT-Q148 and PNKP (lane 5) and analyzed by WBs to measure PNKP levels (blue arrowheads). Mitochondrial protein TFAM used as loading control. Lane 1: Protein molecular weight markers in kDa.

**(D).** DNA isolated from PC12 cells (Cntl; lanes 1 and 2), PC12 cells expressing mHTT-Q148 (lanes 3 and 4) and co-expressing mHTT-Q148 and PNKP (lanes 5 and 6); 13.4 kb and 0.2-kb segments of rat mtDNA PCR amplified, and the PCR products analyzed on agarose gels, stained with EtBr (arrowheads) and analyzed.

**(E).** Relative PCR amplification efficacies of mtDNA segments (in terms of % change) in PC12 cells, and PC12 cells expressing mHTT-Q148, or co-expressing mHTT-Q148 and PNKP. Data represent mean ± SD; ****p<0.0001; ns = not significant.

**(F).** Cell viabilities of PC12 cells expressing wtHTT-Q23, mHTT-Q148 and mutant PC12 cells co-expressing mHTT-Q148 and PNKP measured; Data represent mean ± SD; *p<0.05; **p<0.01; ****p<0.0001.

**(G).** Control PC12 cells, PC12 cells expressing mHTT-Q148 or co-expressing mHTT-Q148 and PNKP treated with JC-10 dye and mitochondrial membrane potential measured. Three biological replicates and five technical replicates were used for this measurement. data represent mean ± SD; **p<0.01; ***p<0.001; ns = not significant.

**Supplementary figure 6. Wildtype HTT expression improves, and in contrast mHTT expression compromises mtDNA integrity in SH-SY5Y cells.**

**(A).** Protein extracts isolated from the control SH-SY5Y cells (Cntl: lane 2), cells expressing wtHTT (wtHTT-Q19 or wtHTT-Q24: lanes 3 and 4), and mHTT with polyQ expansions (mHTT-Q79, mHTT-Q109 and mHTT-Q145; lanes 5 to 7) and HTT levels determined by WBs. TFAM was used as loading control. Lane 1: Protein molecular weight marker in kDa. Proximity ligation assay (PLA) performed to assess relative interaction/ association of HTT with PNKP in control SH-SY5Y cells and cells expressing wtHTT to examine protein interactions. Mitochondria stained with Mito tracker red, and green fluorescence indicates positive protein-protein interactions (arrows show representative positive PLA signals). Nuclei stained with DAPI.

**(B).** PLA with anti-PNKP (rabbit polyclonal NBP1-87257; Novus) and anti-HTT (mouse monoclonal; MAB2170; Millipore-Sigma) Abs in SH-SY5Y cells (Cntl), and SH-SY5Y cells overexpressing wtHTT-Q19 (wtHTT-Q19-OE).

**(C).** PLA with anti-POLRMT (rabbit polyclonal; Genetex GTX105137) and anti-HTT (mouse monoclonal; MAB2170; Millipore-Sigma) Abs in SH-SY5Y cells (Cntl), and SH-SY5Y cells overexpressing wtHTT-Q19 (wtHTT-Q19-OE).

**(D).** Relative PLA signal showing interaction of PNKP with HTT and POLRMT with HTT in control SH-SY5Y cells and in SH-SY5Y cells overexpressing wtHTTQ19. Data represent mean ± SD; **p<0.01.

**(E).** 3’-Phosphatase activities of PNKP in the MEs of control SH-SY5Y cells (Cntl; lane 2), and SH-SY5Y cells expressing wtHTT-Q19 (lane 3), or mHTT-Q145 (lane 4) determined as described above. Lane 1: No protein extract added to the substrate as negative control; Lane 5: Purified PNKP (25 femtomoles) added to the substrate as positive control.

**(F).** Relative 3’-Phosphatase activities (in terms of % product) of PNKP in untreated SH-SY5Y cells (Cntl), and in SH-SY5Y cells expressing wtHTT-Q19 or mHTT-Q145. Three biological and three technical replicates were used for the calculation. Data represent mean ± SD; ****p<0.0001.

**(G).** Total DNA isolated from control SH-SY5Y cells (lanes 1-3), and SH-SY5Y cells ectopically expressing wtHTT-Q19 (lanes 4 to 6) or mHTT-Q145 (lanes 7 to 9), and the 8.9 kb and 0.2 kb segments (arrowheads) of human mtDNA PCR amplified and the PCR products analyzed on agarose gels as described above. Lane 10: 1-kb DNA ladder.

**(H).** Relative PCR amplification efficiencies of mtDNA segments from control SH-SY5Y cells, and in SH-SY5Y cells expressing either wtHTT-Q19 or mHTT-Q145. Three biological and three technical replicates used for the calculation; Data represent mean ± SD; **p<0.01; ***p<0.001; ***p<0.0001.

**(I).** Total DNA isolated from control SH-SY5Y cells (lanes 1 to 3), and SH-SY5Y cells expressing mtHTT-Q145 (lanes 4 to 6) or co-expressing mHTT-Q145 and wtHTT-Q19 (lanes 7 to 9), and the 8.9 kb and 0.2 kb segments of human mtDNA PCR amplified, and the PCR products analyzed on agarose gels as described above (arrowheads). Lane 10: 1-kb DNA ladder.

**(J).** Relative PCR amplification efficiencies of mtDNA segments from control SH-SY5Y cells, and SH-SY5Y cells expressing mHTT, or co-expressing mHTT-Q145 and wtHTT-Q19. Three biological and three technical replicates are used for calculation. Data represents mean ± SD; *p<0.05; **p<0.01; ns = not significant.

**Supplementary figure 7: Transgenic mice expressing the N-terminal truncated fragment of mHTT show reduced mtDNA repair activities and increased mtDNA damage.**

**(A).** PNKP’s 3’-phosphatase activities in MEs isolated from the CRBL (lanes 2 and 3), CTX (lanes 4 and 5), and STR (lanes 6 and 7) of asymptomatic 7-week-old male N171-82Q HD transgenic mice (lanes 3, 5 and 7) and age-matched control (Cntl) mice (lanes 2, 4 and 6); no protein added to the DNA substrate in lane 1 as negative control, and purified PNKP (25 femtomoles) added to the substrate as a positive control in lane 8.

**(B).** Relative PNKP activities in the MEs isolated from the CRBL, CTX, and STR of 7-week-old N171-82Q HD transgenic and age-matched control (Cntl) mice. Three biological replicates and three technical replicates used for this measurement and in D, F, H. Data represent mean ± SD: ****p<0.0001; ns = not significant.

**(C).** Total DNA isolated from the CRBL, CTX, and STR of asymptomatic (7-week-old) N171-82Q HD transgenic and age-matched control (Cntl) mice, 10.06 kb and 0.2 kb segments of mtDNA PCR amplified, and the PCR products electrophoresed on agarose gels The 10.06 kb and 0.2 kb PCR-amplified mtDNA bands shown with arrowheads.

**(D).** Relative PCR amplification efficacies of mtDNA in control (Cntl) and 7-week-old heterozygous asymptomatic N171-82Q transgenic HD mouse brain tissue. Data represent mean ± SD; ****p<0.0001; ns = not significant.

**(E).** MEs isolated from the CRBL, CTX, and STR of asymptomatic (24-week-old) age-matched control (lanes 2, 4, and 6) and R6/2 (lanes 3, 5, and 7) male mice, and 250 ng of the MEs added to the radiolabeled DNA substrate and relative phosphate release from the DNA substrate assessed; no protein added to the DNA substrate as negative control in lane 1, and purified PNKP (25 femtomoles) added to the substrate as a positive control in lane 8. The substrate and released phosphate bands are shown with arrowheads.

**(F).** Relative mitochondrial PNKP activities in the CRBL (lanes 2 vs. 3), CTX (lanes 4 vs. 5), and STR (lanes 6 vs. 7) of control (Cntl) and R6/2 transgenic mice. Data represent mean ± SD; ****p<0.0001; ns = not significant.

**(G).** DNA isolated from the CRBL, CTX, and STR of symptomatic (24-week-old) R6/2 and age-matched control (Cntl) mice. The 10.06 kb and 0.2 kb segments of mtDNA PCR amplified, and the products from control mice (lanes 1, 3, and 5) and R6/2 mice (lanes 2, 4, and 6) were analyzed on agarose gels (arrowheads).

**(H).** Relative PCR amplification efficacy of mtDNA segments in brain tissue from 24-week-old control (Cntl) and R6/2 mice. Data represent mean ± SD; ****p<0.0001. ns = not significant.

## STAR METHODS

### Plasmid constructs, Cell culture, and Plasmid transfection

Construction of plasmids expressing wtHTT-Q23 or mHTT-Q148 was described previously[92]. Plasmid pBacMam2-DiEx-LIC-C-flag_huntingtin_full-length_Q24 (Addgene plasmid # 111742; http://n2t.net/addgene : Addgene _111742), pBacMam2-DiEx-LIC-C-flag_huntingtin_full-length_Q19 (Addgene plasmid # 111741; http://n2t.net/addgene: 111741; RRID:Addgene_111741), pBacMam2-DiEx-LIC-C-Flag_huntingtin_full-length_Q79 (Addgene plasmid# 111729; http://n2t.net/addgene: 111729; RRID: Addgene_111729), pBacMam2-DiEx-LIC-C-flag_huntingtin_full-length_Q109, (Addgene plasmid # 111730; http://n2t.net/addgene: 111730; RRIDAddgene_111730, pBacMam2-DiEx-LIC-C-flag_huntingtin_full-length_Q109, (Addgene plasmid # 111730; http://n2t.net/addgene: 111730; RRID: Addgene_111730), and pBacMam2-DiEx-LIC-C-C-flag_huntingtin_full-length_Q145 (Addgene plasmid#111731; http://n2t.net/addgene: 111731; RRID: Addgene_111731 were gifts from Cheryl Arrowsmith. The full-length huntingtin cDNA was PCR-amplified from these plasmids and sub-cloned into plasmid pcDNA3.1 (Invitrogen, USA) using appropriate DNA linkers to construct plasmids pRGS-wtHTT-Q19, pRGS-wtHTT-Q24, pRGD-mHTT-Q79, pRSG-mHTT-Q109 and pRGS-mHTT-Q145, expressing full-length FLAG-tagged HTT. Plasmids pBiFC-VC155 (Addgene plasmid# 22011; http://n2t.net/addgene: addgene: 22011; RRID: Addgene_22011) and pBiFC-VN173 (Addgene plasmid # 22010; http://n2t.net/addgene: 22010; RRID: Addgene_22010 were gifts from Chang-Deng Hu. The cDNA of human HTT was cloned in-frame into plasmid pBiFC-VN173, and cDNA of POLRMT or POLGA were cloned in-frame into plasmid pBiFC-VC-155 to construct plasmid p-FLAG-VN-HTT and p-HA-VC-POLRMT or p-HA-VC-POLGA respectively. Plasmids carrying the cDNA for POLRMT, TFAM and POLGA were purchased from Origene, USA, and the cDNA of POLRMT, POLGA or TFAM were PCR-amplified with appropriate primers and sub-cloned into plasmid pCMV-Myc-N (Clontech, USA) to construct plasmids p-Myc-POLRMT, p-Myc-POLGA and p-Myc-TFAM expressing Myc-tagged proteins. The PNKP and HTT cDNA were PCR-amplified and sub-cloned into plasmid pSELECT-Puro-mcs (InvivoGen, USA) with appropriate linkers to construct pRGS-PNKP and pRGS-HTT-Q24, respectively. These plasmids were transfected into PC12 or SH-SY5Y cells expressing mHTT-Q148, and the stable clones were selected for puromycin resistance, and PNKP or HTT expressions in these cells were verified by western blots. Plasmids expressing the micro-RNA-adapted shRNA sequences targeting human HTT or PNKP were purchased from Invitrogen, Thermo Fisher, USA.

Human embryonic kidney 293 (HEK293) cells were purchased from Coriell Institute, USA (Cat # CRL-1573) and cultured in EMEM, containing 10% FBS. Human neuroblastoma SH-SY5Y cells were purchased from ATCC, USA (Cat # CRL-2266) and cultured in DMEM containing 15% FBS, in CO_2_ incubator at 37°C. Mouse striatal cells STHdhQ7 and STHdhQ111[36] were obtained from the Coriell Institute, USA, and cultured in DMEM containing 15% FBS, 1% Pen-Strep, and in cell culture incubator at 33°C containing 5% CO_2_. Plasmids encoding the human micro-RNA-adapted shRNA sequences targeting HTT, POLRMT or PNKP (Horizon Discovery, USA) were transfected into SH-SY5Y cells, and the transfected cells were selected for puromycin resistance. The knockdown efficiencies in the shRNA-expressing stable SH-SY5Y cells were assessed by western blots using appropriate antibodies. Plasmids expressing the full-length FLAG-tagged wildtype and mutant HTT cDNA (pRGS-wtHTT-Q19, pRGS-wtHTT-Q24, pRGS-mHTT-Q79, pRGS-mHTT-Q109 and pRGS-mHTT-Q145) were linearized with PvuI and transfected into SH-SY5Y cells with Lipofectamine 2000 reagent (Invitrogen, USA), and the positive transfected cells were selected for G418 resistance, and the stable cells expressing the wtHTT or mHTT transgene were cultured in DMEM containing 15% FBS in CO2 incubator, and transgene expression was assessed by western blot with anti-FLAG or HTT antibodies. Development of rat neuronal PC12 cells carrying the full-length HTT (wtHTT-Q23, mHTT-Q148) was described previously[21, 92] and were cultured in DMEM containing 10% fetal horse serum, 5% fetal bovine serum, 1% penicillin/streptomycin and doxycycline (500 ng/mL). Expression of HTT transgene was induced by withdrawing doxycycline from the media for 5-7 days, and transgene expression was verified by western blots with anti-HTT antibody. SH-SY5Y cells were transfected with plasmids pRGS-wtHTT-Q24 or pRGS-PNKP, encoding HTT and PNKP respectively using Lipofectamine 2000 reagent (Invitrogen, USA) and the transfected cells were selected in medium containing puromycin. Plasmid pRGS-wtHTT-Q24 was linearized with PvuI and transfected into STHdhQ111 cells, and the positive cells were selected for puromycin resistance. All the cell lines used in this study were authenticated by short tandem repeat analysis in the UTMB Molecular Genomics Core. The possible mycoplasma contaminations in all cell lines were routinely performed using GeM Mycoplasma Detection Kit (SIGMA, USA, Cat # MP0025) and cells were found to be free from mycoplasma contamination.

### Preparation of nuclear, cytosolic, and mitochondrial protein extracts

The whole-cell lysate from cell lines was prepared using the RIPA Buffer (Thermo-Fisher Scientific, Cat# 89901). The cytoplasmic and nuclear fractions were prepared using a NE-PER nuclear protein extraction kit (Thermo Scientific, Cat# 78835) following the manufacturer’s instructions. Mitochondria were isolated and purified using the mitochondrial extraction kit (Thermo Scientific, Cat# 89874), and/or following the mitochondria isolation protocol[93–95]. Briefly, approximately 250 mg cortex tissue from freshly sacrificed wildtype mice was harvested, sliced into small pieces, collected in a sterile, pre-chilled centrifuge tube, and the minced tissue was washed with IBc buffer. To prepare 100 ml of IBc buffer, 10 ml of 0.1M Tris–MOPS and 1 ml of EGTA/Tris was added to 20 ml of 1 M sterile sucrose. The volume was adjusted to 100 ml with sterile water and the pH was adjusted to 7.4. The tissue was then mechanically homogenized using a plastic pestle (∼20 strokes) with freshly added IBc buffer at 1:5 ratio [tissue: buffer (w/v)] in ice to obtain a single-cell slurry (monitored under a microscope to ensure cell dissociation). The homogenate was kept in ice for 15 min following pipetting approximately 25-30 times to completely lyse the cells. Following this procedure, the homogenate was centrifuged at 600g for 10 min at 4°C and the supernatant was collected. The supernatant was centrifuged at 7000g for 10 min at 4°C to obtain the mitochondrial pellet. The mitochondria from various cell lines were isolated using the mitochondrial extraction kit (Thermo Scientific, Cat# 89874). The mitochondrial fraction was briefly treated with trypsin to remove the Mt membrane-associated proteins. The mitochondrial pellet was lysed with mitochondrial extraction buffer [50 mM Tris-HCl pH 7.5; 150 mM NaCl; 1 mM EDTA; 1 mM DTT; 1% Triton X-100; 10% Glycerol; protease inhibitor cocktail] to obtain the final mitochondrial lysate, and the extracted proteins were analyzed by western blots.

### Immuno-electron microscopy

For ultrastructural analysis, a monolayer of SH-SY5Y cells were fixed in situ in a mixture of 2.5% formaldehyde and 0.1% glutaraldehyde in 0.05 M cacodylate buffer pH 7.2 containing 0.01% trinitrophenol and 0.03% CaCl2 for 2 hr at room temperature. After washing in 0.1 M cacodylate buffer, cells were scraped off, pelleted and processed further as a pellet. The pellets were stained *en bloc* with 2% aqueous uranyl acetate, dehydrated in 50% and 75% ethanol and embedded in LR White resin medium grade (Electron Microscopy Sciences [EMS], Hatfield, PA, cat#14381-CA). Ultra-thin sections were cut on Leica EM UC7 ultramicrotome (Leica Microsystems, Buffalo Grove, IL) and collected onto Formvar-carbon coated nickel grids. The grids were incubated in a wet chamber on drops of blocking buffer (0.1% BSA and 0.01 M glycine in 0.05 M tris-buffered saline [TBS]) for 15 min at room temperature. Then HTT primary antibody (MAB 2170; Millipore) was used (1:50) in dilution buffer [1% BSA in 0.05 M TBS] for 1 hr at room temperature and then overnight at 4°C. In the next day, HTT antibody-coated grids were washed 5 times for 3 min each, in blocking buffer and incubated with secondary antibody (goat anti-mouse IgG (H+L) conjugated to 10 nm colloidal gold particles (Electron Microscopy Sciences, Hatfield, PA, cat#25128, code: 110.022), which was diluted 1:20 in dilution buffer for 1 hr. at room temperature. After washing in TBS and distilled water for 5 min, the grids were fixed in 2% aqueous glutaraldehyde for 5 min, washed with distilled water for 5 min 3 times, stained with uranyl acetate for 5 min, washed again with distilled water, lead citrate added for 30 sec, then air dried, and examined in a JEM-1400 (JEOL, Japan) transmission electron microscope at 80 kV. Images were acquired with Orius SC2001 digital camera (Gatan, Pleasanton, CA).

### Analysis of HTT-associated DNA repair proteins by co-immunoprecipitation (co-IP)

Co-IP analyses were performed using mitochondrial extracts of neuronal cells according to our reported protocol[21, 96]. In brief, mitochondria from neuronal cells were isolated using a mitochondria isolation kit or mitochondria isolation protocol (described above). Isolated mitochondria were washed with phosphate-buffered saline (PBS), treated with trypsin (1 mg/ml in PBS for 30 minutes) at room temperature to remove contaminating proteins adhering to the outer mitochondrial surface, and washed extensively with PBS. Mitochondrial protein extracts were prepared with mitochondrial extraction buffer (described above) and then treated with benzonase to remove DNA and RNA to avoid nucleic acid-mediated co-IP. Specific target proteins were immunoprecipitated and the resulting immunocomplexes (ICs) were washed extensively with Tris-buffered saline (TBS, 50 mM Tris-HCl [pH 7.5] and 150 mM KCl) containing 1 mM EDTA, 1% Triton X-100, and 10% glycerol. The immunocomplexes (ICs) were eluted from the beads with a solution of 25 mM Tris-HCl (pH 7.5), 500 mM NaCl or with sample buffer dye (NuPAGE) in addition with 2% β-mercaptoethanol, then analyzed by WB for the presence of interacting protein partners using appropriate antibodies.

### *In situ* proximity ligation assays (PLAs)

Neuronal cells were plated on chamber slides and cultured in DMEM containing 10% FBS for 24 hours. Cells were fixed with 4% paraformaldehyde (PFA), permeabilized with 0.2% Tween-20, and washed with 1× PBS. The fixed cells were incubated with primary antibodies for POLRMT (rabbit polyclonal), PNKP (mouse monoclonal), HTT (mouse monoclonal), CSB (mouse monoclonal) and TFAM (mouse monoclonal). Samples were subjected to PLAs using the Duolink PLA kit (Cat # DUO92008, Sigma). Nuclei were stained with DAPI (4’, 6-diamidino-2-phenylindole), and mitochondria were stained with Mito-tracker Red (Thermo-Fisher-Invitrogen; cat# M7512) and PLA signals were visualized under a confocal microscope at 20× magnification.

### HD iPSC culture and differentiation

All the reagents were purchased from Thermo Fisher Scientific (Waltham, MA USA), unless otherwise specified. Three controls: CS25iCTR18n6, CS71iCTR20n6, CS83iCTR33n1 and three HD: CS87iHD50n7, CS03iHD53n3 and CS09iHD109n1 iPSC lines were derived and cultured as previously described on hESC-qualified Matrigel (HD iPSC Consortium, 2017). Once at 70% confluency, neural induction, and further differentiation of neural progenitors to mature neurons was performed as previously described[38]. After 3 weeks of maturation, medium was removed and cells were washed once with PBS pH 7.4, without Mg^2+^ and Ca^2+^. Cells were washed with 4°C PBS pH 7.4, without Mg^2+^ and Ca^2+^, and scraped using a cell scraper, pipetted into a centrifuge tube, and centrifuged at 250 *x g* for 3 minutes. PBS was removed and samples were flash frozen in liquid nitrogen.

### HD transgenic mice

The HD zQ175 mouse model expresses full-length mHTT from the endogenous mouse HTT promoter, and the mutant human HTT exon 1 carrying expanded CAG sequences was inserted into the mouse HTT locus by homologous recombination[42]. The transgenic mouse line R6/2 or B6CBA-Tg (HDexon1)62Gpb/1J line expressing approximately 1 kb of human HTT exon1 carry approximately 100-115 CAG repeats[55]. The N171-82Q transgenic mouse line [B6C3-Tg (HD82Gln) 81Gschi/J mouse line] expresses the truncated N-terminus of human HTT cDNA with a polyglutamine repeat length of 82 under the control of the mouse prion promoter[54]. The heterozygous mice and control littermates (n = 4-5 pools of 2 animals per genotype) were sacrificed, and fresh brain tissues were used to isolate genomic DNA for PNKP assays and proteins for WB analyses. For immunofluorescence assays, transgenic and control mice were deeply anesthetized and transcardially perfused with sterile PBS followed by 4% PFA in PBS. Brains were postfixed overnight in fixative solution and embedded in OCT. Slides with 4-μm-thick frozen sections were processed for immunostaining. Sacrifice and tissue collection were performed according to standard approved procedures following national guidelines and animal protocols.

### Antibodies and western blot analysis

Western blots were performed according to the standard procedure, and each experiment was performed a minimum of three times to ensure reproducible and statistically significant results. The antibodies for DNA polymerase gamma (POLG) (Cell signaling, Cat # 13609); mouse monoclonal anti-HTT antibody (Cat # MAB 2170) were from Millipore-Sigma, USA. The mouse monoclonal anti-POLRMT (Cat # sc-365082), mouse anti-RNA polymerase II (Cat # sc-56767), anti-TFAM (Cat # sc-166965), anti-CSB (Cat # sc-398022), and anti-TOM20 (Cat # sc-17764) antibodies were purchased from Santa Cruz Biotechnology, USA. Rabbit monoclonal Anti-TFAM antibody from Abcam (Cat# ab 176558). The anti-PNKP rabbit polyclonal antibody (Cat # NBP1-87257) was from Novus Biologicals, USA. Anti-MT-CO1 (Mt cytochrome c oxidase) mouse monoclonal antibody (Cat # ab14705) from Abcam, USA. The rabbit anti-POLRMT antibody was obtained from Genetex (Cat # GTX 105137). The anti-beta-Actin mouse monoclonal antibody was from Cell Signaling (Cat # 3700S). Anti-Myc antibody from Takara (Cat # 631206), and from Invitrogen (Cat# MA1-21316) and anti-Flag antibody from Origene (Cat # TA50011), and from Sigma (cat # F3165). The anti-PNKP mouse monoclonal antibody was a kind gift from Dr. Michael Weinfeld (University of Alberta, Canada).

### Measurement of PNKP 3’-phosphatase activity from mitochondrial extracts

The 3’-phosphatase activity of PNKP in the mitochondrial extract (250-500 ng) of cells/ mouse brains or with purified recombinant His-tagged PNKP (25 femtomoles) was conducted as we described previously[91, 96, 97]. A ^32^P-labeled 3’-phosphate-containing 51-mer oligo substrates with a strand break in the middle (5 pmol) was incubated at 37°C for 15 min in buffer A (25 mM Tris-HCl, pH 7.5, 100 mM NaCl, 5 mM MgCl_2_, 1 mM DTT, 10% glycerol and 0.1 μg/μl acetylated BSA) with 5 pmol of unlabeled (cold) substrate. The reaction was stopped by adding buffer B (80% formamide, 10 mM NaOH) and the reaction products were electrophoresed on a 20% Urea-PAGE to measure the amount of 3’ phosphate release from the radio-labelled substrate. The radioactive bands were visualized in PhosphoImager (GE Healthcare, USA). The data were represented as % of the phosphate release (% product) with the total radiolabeled substrate as 100.

### Measurement of mtDNA damage

Total DNA from cells and brain tissue was extracted using the DNeasy™ Blood & Tissue Kit (Qiagen, Cat # 69504). To minimize possible aerial oxidation during genomic DNA isolation, 2, 2, 6, 6-tetramethylpiperidine-*N*-oxyl was added to all solutions to a final concentration of 100 μM immediately before use. LA-QPCR assays were performed to quantify mtDNA lesions as previously described[21, 31, 91]. Preliminary assays were performed to ensure PCR amplification linearity with respect to the number of cycles and DNA concentration. Since amplification of a small region would be independent of DNA damage, a small DNA fragment (∼200 bp) from the same mtDNA segment was also amplified to normalize amplification of the large fragment[31]. The amplified products were electrophoresed and quantitated with ImageJ software (National Institutes of Health, USA). The extent of mtDNA damage was calculated as we previously described[21].

The following primers were used for the LA-QPCR analyses to assess mtDNA damages in human, mouse, rat, and *Drosophila* samples.

Human (8.9 kb)

Forward: 5’-TTTCATCATGCGGAGATGTTGGATGG-3’

Reverse: 5’-TCTAAGCCTCCTTATTCGAGCCGA-3’

Human (0.2 kb)

Forward: 5’-CCCCACAAACCCCATTACTAAACCCA-3’

Reverse: 5’-TTTCATCATGCGGAGATGTTGGATGG-3’

Mouse (10.02 kb)

Forward: 5′-GCCAGCCTGACCCATAGCCATAATAT-3′

Reverse: 5′-GAGAGATTTTATGGGTGTAATGCGG-3′

Mouse (117 bp)

Forward: 5’-CCCAGCTACTACCATCATTCAAGT-3’

Reverse: 5’-GATGGTTTGGGAGATTGGTTGATG-3’

Rat (13.4 kb)

Forward: 5’-AAAATCCCCGCAAACAATGACCACCC-3’

Reverse: 5’-GGCAATTAAGAGTGGGATGGTCGGTT-3’

Rat (0.2 kb)

Forward: 5’-CCTCCCATTCATTATCGCCGCCCTTGC-3’

Reverse: 5’-GTCTGGGTCTCCTAGTAGGTCTGGGAA-3’

*Drosophila* (14.2 kb)

Forward: 5’-GCCGCTCCTTTCCATTTTTGATTTCC-3’

Reverse: 5’-TGCCAGCAGTCGCGGTTATACCA-3’

*Drosophila* (151 bp)

Forward: 5’-GCTCCTGATATAGCATTCCCACGA-3’

Reverse: 5’-CATGAGCAATTCCAGCGGATAAA-3’

### Chromatin immunoprecipitation (ChIP) and Sequential ChIP (ChIP-re-ChIP) assay

ChIP assays were performed using fresh mouse brain tissue as previously described[98] with minor modifications. Briefly, 80-100 mg freshly harvested cortical tissues were chopped into small pieces (between 1 and 3 mm^3^) using a scalpel razor and fixed in 1% formaldehyde for 15 min with gentle agitation at room temperature to cross-link DNA to bound proteins. The samples were centrifuged at 440 × *g* for 5 min at room temperature, followed by the addition of 0.125 M glycine to terminate the cross-linking reaction. The samples were washed three times with ice-cold PBS (containing a protease inhibitor mixture) and centrifuged each time at 440 × *g* for 5 min at 4°C. The pellet was re-suspended in 1 ml ice-cold lysis buffer (10 mM EDTA, 1% [w/v] SDS, 50 mM Tris-HCl [pH 7.5]) with protease inhibitors for 15 min on ice and homogenized slowly (∼20 strokes on ice) with a hand homogenizer to produce a single-cell suspension. Then, the samples were subjected to sonication for lysis as well as to generate ∼500 bp of DNA fragments (sonicated at 30% amplitude in Qsonica sonicator; 5× [3×30 s for each pulse, 1 min hold between every pulse]. The sonicated samples were centrifuged at 15,000 × *g* for 30 min at 4°C, and the supernatants were collected for ChIP analysis as described previously[98]. After the recovery of DNA with proteinase K treatment followed by phenol extraction and ethanol precipitation, 1% of input chromatin and the precipitated DNA were analyzed by qPCR with the primer sets listed below. The ChIP data are presented as percent binding relative to the input value. The same protocol was followed in the case of adherent cells. Briefly, the mutant and control cells were grown on 10-cm tissue culture plates in DMEM containing 10% FBS. Formaldehyde was added to the medium to a final concentration of 1%, and the cells were incubated for 15 minutes. The reaction was stopped by adding 0.125 M glycine and the same procedure was followed as described above that we used for the mouse tissue. The sheared chromatin was immunoprecipitated for overnight at 4°C with 10 μg isotype control IgG (SC-2025; Santa Cruz Biotechnology, USA) or anti-HTT (Millipore; Cat #MAB2170) or PNKP (Novus biologicals; Cat# NBP1-87257) antibodies. MtDNA sequences were detected with primers specifically for the mitochondrial D-loop, cytochrome c oxidase 1 and 2 (Mt-co1 and Mt-co2 respectively), NADH dehydrogenase 2 (Mt-nd2) and mtDNA-coded cytochrome b (Mt-cytb).

For re-ChIP assays, the primary immunocomplexes were eluted with ChIP elution buffer (0.1M NaHCO3, 1% SDS, protease inhibitor, at 50^0^C for an hour), obtained from either anti-HTT or anti-POLRMT (Genetex; Cat# GTX 105137) antibodies. The eluant was diluted 10-fold with ChIP dilution buffer (20 mM Tris-HCl, pH 8.0, 1 mM EDTA, 150 mM NaCl, 1% Triton X-100, and protease inhibitors) and then subjected to further immunoprecipitation with the anti-HTT or anti POLRMT or control IgG Ab using identical procedure.

The following primers were used to detect mouse mtDNA in ChIP and ChIP-re-ChIP experiments described in this manuscript.

**Mt-co1:**

Forward: 5’-CTGAGCGGGAATAGTGGGTA-3’

Reverse: 5’-TGGGGCTCCGATTATTAGTG-3’

**Mt-co2:**

Forward: 5’-TCTCCCCTCTCTACGCATTC-3’

Reverse: 5’-GGCAGAACGACTCGGTTATC-3’

**Mt-nd2:**

Forward: 5’-CCTTCGACCTGACAGAAGGA-3’

Reverse: 5’-GATGCTCGGATCCATAGGAA-3’

**Mt-Cytb:**

Forward: 5’-ACGTCCTTCCATGAGGACAA-3’

Reverse: 5’-GAGGTGAACGATTGCTAGGG-3’

**D-loop:**

Forward: 5’-TGCGTTATCGCCCATACGTT-3’

Reverse: 5’-AGTGTTTTTGGGGTTTGGCATT-3’

### Mitochondrial membrane potential detection

The mutant and controls cells (PC12 cells expressing either wtHTT-Q23 or mHTT-Q148) overexpressing PNKP were cultured in DMEM supplemented with 10% FBS, 1 penicillin and streptomycin, 2mM L-glutamine, and 44 μg/ml G418. Cells were grown at 33°C in a 5% CO_2_ incubator, then seeded in 10 cm cell culture plates and stained with JC-10. Briefly, 6 × 10^5^ cells were harvested, fixed with 0.5 ml cell culture media and 0.5 ml JC-10 staining solution, and incubated in a cell culture cabinet for 20 min after gentle shaking. Cells were collected by centrifugation at 600 × g, 4°C for 3 min and then washed three times with 1× JC-10 buffer. Cell fluorescence intensity was measured with CytoFLEX flow cytometer (Beckman, USA) after cells were resuspended in 100 μl 1× JC-10 buffer.

### Cell proliferation or viability assay

Cell toxicity in SH-SY5Y cells, SH-SY5Y cells expressing HTT-shRNA or overexpressing HTT cDNA were measured using an MTT assay kit (Cat # ab211091, Abcam) and using the instructions provided by the manufacturer.

### Image collection

The confocal images were collected using a Zeiss LSM-510 META confocal microscope with 40× or 60× 1.2 numerical aperture water immersion objectives. Images were obtained using two excitation wavelengths (488 and 543 nm) by sequential acquisition. Images were collected using 4-frame-Kallman-averaging with a pixel time of 1.26 μs, a pixel size of 110 nm, and optical slices of 1.0 μm. Z-stack acquisition was performed at 0.8-μm steps. Orthogonal views were processed with LSM 510 software.

### HD autopsy brain tissue samples

Human autopsy specimens (caudate nucleus) were obtained from NIH NeuroBiobank, in accordance with local legislation and ethical rules. Control brain samples were collected from age-matched individuals without neurodegenerative disorders. The HD brain tissue samples were obtained from patients with HD who were clinically characterized based on the presence of chorea and motor, mood, and cognitive impairment. The molecular diagnosis of HD was established by analyzing genomic DNA extracted from peripheral blood using a combination of PCR and Southern blotting. The CAG repeat lengths in HTT were established by sequencing the expansion loci of the mutant allele. Human brain samples were dissected, and flash frozen in liquid nitrogen and stored at -80°C until further analysis. Genomic DNA isolated from the caudate of HD patients’ and age-matched control brain tissue (caudate). The HD patients’ brain tissues that are used for the mtDNA damage analyses in this manuscript are: (1) 56 years-old, male; grade 2; (2). 58 years-old, male; grade 2; (3) 53 years-old, male; grade 3; (4) 52 years-old, male; grade 3; (5) 85 years-old, male; grade 3; (6) 53 years-old, female; grade 3. The 10.06 and 0.2 kb sections of mtDNA PCR amplified, and the PCR products from control and HD analyzed on agarose gels and the DNA bands quantified. Five biological replicates and three technical replicates were used in this study.

### *Drosophila* Climbing Assay to assess motor function in *Drosophila*

Age matched *Drosophila* from WT, WT flies expressing full-length HTT with 16 CAG repeats (HTT-16Q) and WT flies expressing mutant huntingtin with 128 CAG repeats (HTT-128Q) were raised at 25°C for 20 days. On the 21^st^ day, flies from each genotype were used individually to test for their motor function using the climbing assay method described[63] (with minor changes. As such, flies from each genotype were transferred to a clear plastic cylinder with the climbing line to cross being marked at 12.5cm. The number of flies crossing the line was recorded every 5 seconds and the quantification was performed using two-way ANOVA between the individual time points for all genotypes. A total of 25 individual flies were used per genotype and the experiments were repeated 4 times.

### Mitochondrial HTT complex objective computational modeling

Due to the large size of HTT (∼350 kDa), alphafold models (HTT-POLGA, HTT-PNKP, HTT-TFAM) were predicted separately using AlphaFold3 server (https://alphafoldserver.com/). Predicted local distance difference test (pLDDT) confidence score is shown in Figure 2G. The overall predicted template modeling (pTM) scores range between 0.61 and 0.69. A pTM score above 0.5 indicates that the overall predicted fold for the complex may be comparable to the true structure. Because polyQ expansion affects PNKP SUMOylation and activity based on [21], the model with PNKP near the polyQ area was selected, results in the only non-clash HTT-POLGA-PNKP model out of 5 predicted models. TFAM is also consistently predicted to bind on the N-HEAT domain of HTT with minimal clash with POLGA.

### Statistical analysis

Data reported as mean ± SD and the statistical analysis was performed using GraphPad Prism software. Differences between two experimental groups were analyzed by Student’s t test (2-tail, assuming unequal variances). When comparing multiple groups, One-way ANOVA was performed followed by Holm-Sidak test to determine significance. Statistical analyses of all data were performed using a t test in GraphPad Prism Version 7.03 (*P < 0.005, ** P < 0.01, *** P < 0.001, **** P < 0.0001).

## REFERENCES

1. Galimberti M, Nucera MR, Bocchi VD, Conforti P, Vezzoli E, Cereda M, Maffezzini C, Iennaco R, Scolz A, Falqui A, et al: Huntington’s disease cellular phenotypes are rescued non-cell autonomously by healthy cells in mosaic telencephalic organoids. Nat Commun 2024, 15:6534.

2. Ho LW, Brown R, Maxwell M, Wyttenbach A, Rubinsztein DC: Wild type Huntingtin reduces the cellular toxicity of mutant Huntingtin in mammalian cell models of Huntington’s disease. J Med Genet 2001, 38:450–452.

3. Leavitt BR, van Raamsdonk JM, Shehadeh J, Fernandes H, Murphy Z, Graham RK, Wellington CL, Raymond LA, Hayden MR: Wild-type huntingtin protects neurons from excitotoxicity. J Neurochem 2006, 96:1121–1129.

4. Rigamonti D, Bauer JH, De-Fraja C, Conti L, Sipione S, Sciorati C, Clementi E, Hackam A, Hayden MR, Li Y, et al: Wild-type huntingtin protects from apoptosis upstream of caspase-3. J Neurosci 2000, 20:3705–3713.

5. Zhang Y, Leavitt BR, van Raamsdonk JM, Dragatsis I, Goldowitz D, MacDonald ME, Hayden MR, Friedlander RM: Huntingtin inhibits caspase-3 activation. EMBO J 2006, 25:5896–5906.

6. Zhang Y, Li M, Drozda M, Chen M, Ren S, Mejia Sanchez RO, Leavitt BR, Cattaneo E, Ferrante RJ, Hayden MR, Friedlander RM: Depletion of wild-type huntingtin in mouse models of neurologic diseases. J Neurochem 2003, 87:101–106.

7. Dragatsis I, Levine MS, Zeitlin S: Inactivation of Hdh in the brain and testis results in progressive neurodegeneration and sterility in mice. Nat Genet 2000, 26:300–306.

8. Ismailoglu I, Chen Q, Popowski M, Yang L, Gross SS, Brivanlou AH: Huntingtin protein is essential for mitochondrial metabolism, bioenergetics and structure in murine embryonic stem cells. Dev Biol 2014, 391:230–240.

9. A novel gene containing a trinucleotide repeat that is expanded and unstable on Huntington’s disease chromosomes. The Huntington’s Disease Collaborative Research Group. Cell 1993, 72:971–983.

10. Bossy-Wetzel E, Petrilli A, Knott AB: Mutant huntingtin and mitochondrial dysfunction. Trends Neurosci 2008, 31:609–616.

11. Choo YS, Johnson GV, MacDonald M, Detloff PJ, Lesort M: Mutant huntingtin directly increases susceptibility of mitochondria to the calcium-induced permeability transition and cytochrome c release. Hum Mol Genet 2004, 13:1407– 1420.

12. Panov AV, Gutekunst CA, Leavitt BR, Hayden MR, Burke JR, Strittmatter WJ, Greenamyre JT: Early mitochondrial calcium defects in Huntington’s disease are a direct effect of polyglutamines. Nat Neurosci 2002, 5:731–736.

13. Song W, Chen J, Petrilli A, Liot G, Klinglmayr E, Zhou Y, Poquiz P, Tjong J, Pouladi MA, Hayden MR, et al: Mutant huntingtin binds the mitochondrial fission GTPase dynamin-related protein-1 and increases its enzymatic activity. Nat Med 2011, 17:377–382.

14. Acevedo-Torres K, Berrios L, Rosario N, Dufault V, Skatchkov S, Eaton MJ, Torres-Ramos CA, Ayala-Torres S: Mitochondrial DNA damage is a hallmark of chemically induced and the R6/2 transgenic model of Huntington’s disease. DNA Repair (Amst) 2009, 8:126–136.

15. Ayala-Pena S: Role of oxidative DNA damage in mitochondrial dysfunction and Huntington’s disease pathogenesis. Free Radic Biol Med 2013, 62:102–110.

16. Browne SE, Bowling AC, MacGarvey U, Baik MJ, Berger SC, Muqit MM, Bird ED, Beal MF: Oxidative damage and metabolic dysfunction in Huntington’s disease: selective vulnerability of the basal ganglia. Ann Neurol 1997, 41:646–653.

17. Polidori MC, Mecocci P, Browne SE, Senin U, Beal MF: Oxidative damage to mitochondrial DNA in Huntington’s disease parietal cortex. Neurosci Lett 1999, 272:53–56.

18. Siddiqui A, Rivera-Sanchez S, Castro Mdel R, Acevedo-Torres K, Rane A, Torres-Ramos CA, Nicholls DG, Andersen JK, Ayala-Torres S: Mitochondrial DNA damage is associated with reduced mitochondrial bioenergetics in Huntington’s disease. Free Radic Biol Med 2012, 53:1478–1488.

19. Maiuri T, Mocle AJ, Hung CL, Xia J, van Roon-Mom WM, Truant R: Huntingtin is a scaffolding protein in the ATM oxidative DNA damage response complex. Hum Mol Genet 2017, 26:395–406.

20. Ochaba J, Lukacsovich T, Csikos G, Zheng S, Margulis J, Salazar L, Mao K, Lau AL, Yeung SY, Humbert S, et al: Potential function for the Huntingtin protein as a scaffold for selective autophagy. Proc Natl Acad Sci U S A 2014, 111:16889–16894.

21. Gao R, Chakraborty A, Geater C, Pradhan S, Gordon KL, Snowden J, Yuan S, Dickey AS, Choudhary S, Ashizawa T, et al: Mutant huntingtin impairs PNKP and ATXN3, disrupting DNA repair and transcription. Elife 2019, 8.

22. Shokolenko I, Venediktova N, Bochkareva A, Wilson GL, Alexeyev MF: Oxidative stress induces degradation of mitochondrial DNA. Nucleic Acids Res 2009, 37:2539– 2548.

23. Weissman L, de Souza-Pinto NC, Stevnsner T, Bohr VA: DNA repair, mitochondria, and neurodegeneration. Neuroscience 2007, 145:1318–1329.

24. Shirendeb U, Reddy AP, Manczak M, Calkins MJ, Mao P, Tagle DA, Reddy PH: Abnormal mitochondrial dynamics, mitochondrial loss and mutant huntingtin oligomers in Huntington’s disease: implications for selective neuronal damage. Hum Mol Genet 2011, 20:1438–1455.

25. Chakraborty J, Rajamma U, Mohanakumar KP: A mitochondrial basis for Huntington’s disease: therapeutic prospects. Mol Cell Biochem 2014, 389:277–291.

26. Sonsky I, Vodicka P, Vodickova Kepkova K, Hansikova H: Mitophagy in Huntington’s disease. Neurochem Int 2021, 149:105147.

27. Mandal SM, Hegde ML, Chatterjee A, Hegde PM, Szczesny B, Banerjee D, Boldogh I, Gao R, Falkenberg M, Gustafsson CM, et al: Role of human DNA glycosylase Nei-like 2 (NEIL2) and single strand break repair protein polynucleotide kinase 3’-phosphatase in maintenance of mitochondrial genome. J Biol Chem 2012, 287:2819– 2829.

28. Aamann MD, Sorensen MM, Hvitby C, Berquist BR, Muftuoglu M, Tian J, de Souza-Pinto NC, Scheibye-Knudsen M, Wilson DM, 3rd, Stevnsner T, Bohr VA: Cockayne syndrome group B protein promotes mitochondrial DNA stability by supporting the DNA repair association with the mitochondrial membrane. FASEB J 2010, 24:2334– 2346.

29. Kamenisch Y, Fousteri M, Knoch J, von Thaler AK, Fehrenbacher B, Kato H, Becker T, Dolle ME, Kuiper R, Majora M, et al: Proteins of nucleotide and base excision repair pathways interact in mitochondria to protect from loss of subcutaneous fat, a hallmark of aging. J Exp Med 2010, 207:379–390.

30. Salazar JJ, Van Houten B: Preferential mitochondrial DNA injury caused by glucose oxidase as a steady generator of hydrogen peroxide in human fibroblasts. Mutat Res 1997, 385:139–149.

31. Santos JH, Meyer JN, Mandavilli BS, Van Houten B: Quantitative PCR-based measurement of nuclear and mitochondrial DNA damage and repair in mammalian cells. Methods Mol Biol 2006, 314:183–199.

32. Boetefuer EL, Lake RJ, Fan HY: Mechanistic insights into the regulation of transcription and transcription-coupled DNA repair by Cockayne syndrome protein B. Nucleic Acids Res 2018, 46:7471–7479.

33. Menoni H, Wienholz F, Theil AF, Janssens RC, Lans H, Campalans A, Radicella JP, Marteijn JA, Vermeulen W: The transcription-coupled DNA repair-initiating protein CSB promotes XRCC1 recruitment to oxidative DNA damage. Nucleic Acids Res 2018, 46:7747–7756.

34. van den Boom V, Citterio E, Hoogstraten D, Zotter A, Egly JM, van Cappellen WA, Hoeijmakers JH, Houtsmuller AB, Vermeulen W: DNA damage stabilizes interaction of CSB with the transcription elongation machinery. J Cell Biol 2004, 166:27–36.

35. Abramson J, Adler J, Dunger J, Evans R, Green T, Pritzel A, Ronneberger O, Willmore L, Ballard AJ, Bambrick J, et al: Accurate structure prediction of biomolecular interactions with AlphaFold 3. Nature 2024, 630:493–500.

36. Trettel F, Rigamonti D, Hilditch-Maguire P, Wheeler VC, Sharp AH, Persichetti F, Cattaneo E, MacDonald ME: Dominant phenotypes produced by the HD mutation in STHdh(Q111) striatal cells. Hum Mol Genet 2000, 9:2799–2809.

37. Furlan-Magaril M, Rincon-Arano H, Recillas-Targa F: Sequential chromatin immunoprecipitation protocol: ChIP-reChIP. Methods Mol Biol 2009, 543:253–266.

38. Smith-Geater C, Hernandez SJ, Lim RG, Adam M, Wu J, Stocksdale JT, Wassie BT, Gold MP, Wang KQ, Miramontes R, et al: Aberrant Development Corrected in Adult-Onset Huntington’s Disease iPSC-Derived Neuronal Cultures via WNT Signaling Modulation. Stem Cell Reports 2020, 14:406–419.

39. Madabhushi R, Pan L, Tsai LH: DNA damage and its links to neurodegeneration. Neuron 2014, 83:266–282.

40. Ross CA, Truant R: DNA repair: A unifying mechanism in neurodegeneration. Nature 2017, 541:34–35.

41. Shanbhag NM, Evans MD, Mao W, Nana AL, Seeley WW, Adame A, Rissman RA, Masliah E, Mucke L: Early neuronal accumulation of DNA double strand breaks in Alzheimer’s disease. Acta Neuropathol Commun 2019, 7:77.

42. Menalled LB, Kudwa AE, Miller S, Fitzpatrick J, Watson-Johnson J, Keating N, Ruiz M, Mushlin R, Alosio W, McConnell K, et al: Comprehensive behavioral and molecular characterization of a new knock-in mouse model of Huntington’s disease: zQ175. PLoS One 2012, 7:e49838.

43. Vonsattel JP, DiFiglia M: Huntington disease. J Neuropathol Exp Neurol 1998, 57:369– 384.

44. Hewitt VL, Whitworth AJ: Mechanisms of Parkinson’s Disease: Lessons from Drosophila. Curr Top Dev Biol 2017, 121:173–200.

45. Krench M, Littleton JT: Neurotoxicity Pathways in Drosophila Models of the Polyglutamine Disorders. Curr Top Dev Biol 2017, 121:201–223.

46. Marsh JL, Pallos J, Thompson LM: Fly models of Huntington’s disease. Hum Mol Genet 2003, 12 Spec No 2:R187–193.

47. Wentzell J, Kretzschmar D: Alzheimer’s disease and tauopathy studies in flies and worms. Neurobiol Dis 2010, 40:21–28.

48. Rosas-Arellano A, Estrada-Mondragon A, Pina R, Mantellero CA, Castro MA: The Tiny Drosophila Melanogaster for the Biggest Answers in Huntington’s Disease. Int J Mol Sci 2018, 19.

49. Weiss KR, Littleton JT: Characterization of axonal transport defects in Drosophila Huntingtin mutants. J Neurogenet 2016, 30:212–221.

50. Sathasivam K, Neueder A, Gipson TA, Landles C, Benjamin AC, Bondulich MK, Smith DL, Faull RL, Roos RA, Howland D, et al: Aberrant splicing of HTT generates the pathogenic exon 1 protein in Huntington disease. Proc Natl Acad Sci U S A 2013, 110:2366–2370.

51. Sawa A, Nagata E, Sutcliffe S, Dulloor P, Cascio MB, Ozeki Y, Roy S, Ross CA, Snyder SH: Huntingtin is cleaved by caspases in the cytoplasm and translocated to the nucleus via perinuclear sites in Huntington’s disease patient lymphoblasts. Neurobiol Dis 2005, 20:267–274.

52. Wellington CL, Ellerby LM, Gutekunst CA, Rogers D, Warby S, Graham RK, Loubser O, van Raamsdonk J, Singaraja R, Yang YZ, et al: Caspase cleavage of mutant huntingtin precedes neurodegeneration in Huntington’s disease. J Neurosci 2002, 22:7862–7872.

53. Orr AL, Li S, Wang CE, Li H, Wang J, Rong J, Xu X, Mastroberardino PG, Greenamyre JT, Li XJ: N-terminal mutant huntingtin associates with mitochondria and impairs mitochondrial trafficking. J Neurosci 2008, 28:2783–2792.

54. Schilling G, Becher MW, Sharp AH, Jinnah HA, Duan K, Kotzuk JA, Slunt HH, Ratovitski T, Cooper JK, Jenkins NA, et al: Intranuclear inclusions and neuritic aggregates in transgenic mice expressing a mutant N-terminal fragment of huntingtin. Hum Mol Genet 1999, 8:397–407.

55. Mangiarini L, Sathasivam K, Seller M, Cozens B, Harper A, Hetherington C, Lawton M, Trottier Y, Lehrach H, Davies SW, Bates GP: Exon 1 of the HD gene with an expanded CAG repeat is sufficient to cause a progressive neurological phenotype in transgenic mice. Cell 1996, 87:493–506.

56. Seong IS, Ivanova E, Lee JM, Choo YS, Fossale E, Anderson M, Gusella JF, Laramie JM, Myers RH, Lesort M, MacDonald ME: HD CAG repeat implicates a dominant property of huntingtin in mitochondrial energy metabolism. Hum Mol Genet 2005, 14:2871–2880.

57. Weydt P, Pineda VV, Torrence AE, Libby RT, Satterfield TF, Lazarowski ER, Gilbert ML, Morton GJ, Bammler TK, Strand AD, et al: Thermoregulatory and metabolic defects in Huntington’s disease transgenic mice implicate PGC-1alpha in Huntington’s disease neurodegeneration. Cell Metab 2006, 4:349–362.

58. Brand AH, Perrimon N: Targeted gene expression as a means of altering cell fates and generating dominant phenotypes. Development 1993, 118:401–415.

59. Budnik V, Koh YH, Guan B, Hartmann B, Hough C, Woods D, Gorczyca M: Regulation of synapse structure and function by the Drosophila tumor suppressor gene dlg. Neuron 1996, 17:627–640.

60. Caviston JP, Holzbaur EL: Huntingtin as an essential integrator of intracellular vesicular trafficking. Trends Cell Biol 2009, 19:147–155.

61. Caviston JP, Ross JL, Antony SM, Tokito M, Holzbaur EL: Huntingtin facilitates dynein/dynactin-mediated vesicle transport. Proc Natl Acad Sci U S A 2007, 104:10045–10050.

62. Saudou F, Humbert S: The Biology of Huntingtin. Neuron 2016, 89:910–926.

63. Madabattula ST, Strautman JC, Bysice AM, O’Sullivan JA, Androschuk A, Rosenfelt C, Doucet K, Rouleau G, Bolduc F: Quantitative Analysis of Climbing Defects in a Drosophila Model of Neurodegenerative Disorders. J Vis Exp 2015:e52741.

64. Horton TM, Graham BH, Corral-Debrinski M, Shoffner JM, Kaufman AE, Beal MF, Wallace DC: Marked increase in mitochondrial DNA deletion levels in the cerebral cortex of Huntington’s disease patients. Neurology 1995, 45:1879–1883.

65. Mochel F, Haller RG: Energy deficit in Huntington disease: why it matters. J Clin Invest 2011, 121:493–499.

66. Tellez-Nagel I, Johnson AB, Terry RD: Studies on brain biopsies of patients with Huntington’s chorea. J Neuropathol Exp Neurol 1974, 33:308–332.

67. Djousse L, Knowlton B, Cupples LA, Marder K, Shoulson I, Myers RH: Weight loss in early stage of Huntington’s disease. Neurology 2002, 59:1325–1330.

68. Gu M, Gash MT, Mann VM, Javoy-Agid F, Cooper JM, Schapira AH: Mitochondrial defect in Huntington’s disease caudate nucleus. Ann Neurol 1996, 39:385–389.

69. Aidt FH, Nielsen SM, Kanters J, Pesta D, Nielsen TT, Norremolle A, Hasholt L, Christiansen M, Hagen CM: Dysfunctional mitochondrial respiration in the striatum of the Huntington’s disease transgenic R6/2 mouse model. PLoS Curr 2013, 5.

70. Gines S, Seong IS, Fossale E, Ivanova E, Trettel F, Gusella JF, Wheeler VC, Persichetti F, MacDonald ME: Specific progressive cAMP reduction implicates energy deficit in presymptomatic Huntington’s disease knock-in mice. Hum Mol Genet 2003, 12:497– 508.

71. Milakovic T, Johnson GV: Mitochondrial respiration and ATP production are significantly impaired in striatal cells expressing mutant huntingtin. J Biol Chem 2005, 280:30773–30782.

72. Tabrizi SJ, Workman J, Hart PE, Mangiarini L, Mahal A, Bates G, Cooper JM, Schapira AH: Mitochondrial dysfunction and free radical damage in the Huntington R6/2 transgenic mouse. Ann Neurol 2000, 47:80–86.

73. Brosey CA, Shen R, Tainer JA: NADH-bound AIF activates the mitochondrial CHCHD4/MIA40 chaperone by a substrate-mimicry mechanism. EMBO J 2025, 44:1220–1248.

74. Eckelmann BJ, Bacolla A, Wang H, Ye Z, Guerrero EN, Jiang W, El-Zein R, Hegde ML, Tomkinson AE, Tainer JA, Mitra S: XRCC1 promotes replication restart, nascent fork degradation and mutagenic DNA repair in BRCA2-deficient cells. NAR Cancer 2020, 2:zcaa013.

75. Hambarde S, Tsai CL, Pandita RK, Bacolla A, Maitra A, Charaka V, Hunt CR, Kumar R, Limbo O, Le Meur R, et al: EXO5-DNA structure and BLM interactions direct DNA resection critical for ATR-dependent replication restart. Mol Cell 2021, 81:2989– 3006 e2989.

76. Sparks JL, Gerik KJ, Stith CM, Yoder BL, Burgers PM: The roles of fission yeast exonuclease 5 in nuclear and mitochondrial genome stability. DNA Repair (Amst) 2019, 83:102720.

77. Ye Z, Shi Y, Lees-Miller SP, Tainer JA: Function and Molecular Mechanism of the DNA Damage Response in Immunity and Cancer Immunotherapy. Front Immunol 2021, 12:797880.

78. Hilditch-Maguire P, Trettel F, Passani LA, Auerbach A, Persichetti F, MacDonald ME: Huntingtin: an iron-regulated protein essential for normal nuclear and perinuclear organelles. Hum Mol Genet 2000, 9:2789–2797.

79. Chatterjee A, Saha S, Chakraborty A, Silva-Fernandes A, Mandal SM, Neves-Carvalho A, Liu Y, Pandita RK, Hegde ML, Hegde PM, et al: The role of the mammalian DNA end-processing enzyme polynucleotide kinase 3’-phosphatase in spinocerebellar ataxia type 3 pathogenesis. PLoS Genet 2015, 11:e1004749.

80. Gao R, Liu Y, Silva-Fernandes A, Fang X, Paulucci-Holthauzen A, Chatterjee A, Zhang HL, Matsuura T, Choudhary S, Ashizawa T, et al: Inactivation of PNKP by mutant ATXN3 triggers apoptosis by activating the DNA damage-response pathway in SCA3. PLoS Genet 2015, 11:e1004834.

81. Karimi-Busheri F, Daly G, Robins P, Canas B, Pappin DJ, Sgouros J, Miller GG, Fakhrai H, Davis EM, Le Beau MM, Weinfeld M: Molecular characterization of a human DNA kinase. J Biol Chem 1999, 274:24187–24194.

82. Yakes FM, Van Houten B: Mitochondrial DNA damage is more extensive and persists longer than nuclear DNA damage in human cells following oxidative stress. Proc Natl Acad Sci U S A 1997, 94:514–519.

83. Rockabrand E, Slepko N, Pantalone A, Nukala VN, Kazantsev A, Marsh JL, Sullivan PG, Steffan JS, Sensi SL, Thompson LM: The first 17 amino acids of Huntingtin modulate its sub-cellular localization, aggregation and effects on calcium homeostasis. Hum Mol Genet 2007, 16:61–77.

84. Yu ZX, Li SH, Evans J, Pillarisetti A, Li H, Li XJ: Mutant huntingtin causes context-dependent neurodegeneration in mice with Huntington’s disease. J Neurosci 2003, 23:2193–2202.

85. Gutekunst CA, Li SH, Yi H, Ferrante RJ, Li XJ, Hersch SM: The cellular and subcellular localization of huntingtin-associated protein 1 (HAP1): comparison with huntingtin in rat and human. J Neurosci 1998, 18:7674–7686.

86. Petrasch-Parwez E, Nguyen HP, Lobbecke-Schumacher M, Habbes HW, Wieczorek S, Riess O, Andres KH, Dermietzel R, Von Horsten S: Cellular and subcellular localization of Huntingtin [corrected] aggregates in the brain of a rat transgenic for Huntington disease. J Comp Neurol 2007, 501:716–730.

87. Hanawalt PC, Spivak G: Transcription-coupled DNA repair: two decades of progress and surprises. Nat Rev Mol Cell Biol 2008, 9:958–970.

88. Youle RJ, Narendra DP: Mechanisms of mitophagy. Nat Rev Mol Cell Biol 2011, 12:9– 14.

89. Tabrizi SJ, Ghosh R, Leavitt BR: Huntingtin Lowering Strategies for Disease Modification in Huntington’s Disease. Neuron 2019, 101:801–819.

90. Kwon D: Failure of genetic therapies for Huntington’s devastates community. Nature 2021, 593:180.

91. Morozko EL, Smith-Geater C, Monteys AM, Pradhan S, Lim RG, Langfelder P, Kachemov M, Kulkarni JA, Zaifman J, Hill A, et al: PIAS1 modulates striatal transcription, DNA damage repair, and SUMOylation with relevance to Huntington’s disease. Proc Natl Acad Sci U S A 2021, 118.

92. Igarashi S, Morita H, Bennett KM, Tanaka Y, Engelender S, Peters MF, Cooper JK, Wood JD, Sawa A, Ross CA: Inducible PC12 cell model of Huntington’s disease shows toxicity and decreased histone acetylation. Neuroreport 2003, 14:565–568.

93. Clayton DA, Shadel GS: Isolation of mitochondria from cells and tissues. Cold Spring Harb Protoc 2014, 2014:pdb top074542.

94. Frezza C, Cipolat S, Scorrano L: Organelle isolation: functional mitochondria from mouse liver, muscle and cultured fibroblasts. Nat Protoc 2007, 2:287–295.

95. Liao PC, Bergamini C, Fato R, Pon LA, Pallotti F: Isolation of mitochondria from cells and tissues. Methods Cell Biol 2020, 155:3–31.

96. Chakraborty A, Tapryal N, Venkova T, Mitra J, Vasquez V, Sarker AH, Duarte-Silva S, Huai W, Ashizawa T, Ghosh G, et al: Deficiency in classical nonhomologous end-joining-mediated repair of transcribed genes is linked to SCA3 pathogenesis. Proc Natl Acad Sci U S A 2020, 117:8154–8165.

97. Wiederhold L, Leppard JB, Kedar P, Karimi-Busheri F, Rasouli-Nia A, Weinfeld M, Tomkinson AE, Izumi T, Prasad R, Wilson SH, et al: AP endonuclease-independent DNA base excision repair in human cells. Mol Cell 2004, 15:209–220.

98. Chakraborty A, Tapryal N, Venkova T, Horikoshi N, Pandita RK, Sarker AH, Sarkar PS, Pandita TK, Hazra TK: Classical non-homologous end-joining pathway utilizes nascent RNA for error-free double-strand break repair of transcribed genes. Nat Commun 2016, 7:13049.

