## Supplementary figures and images for "Huntingtin preserves mitochondrial genome integrity in neurons, which is impaired in Huntington’s disease"

### Supplemental data

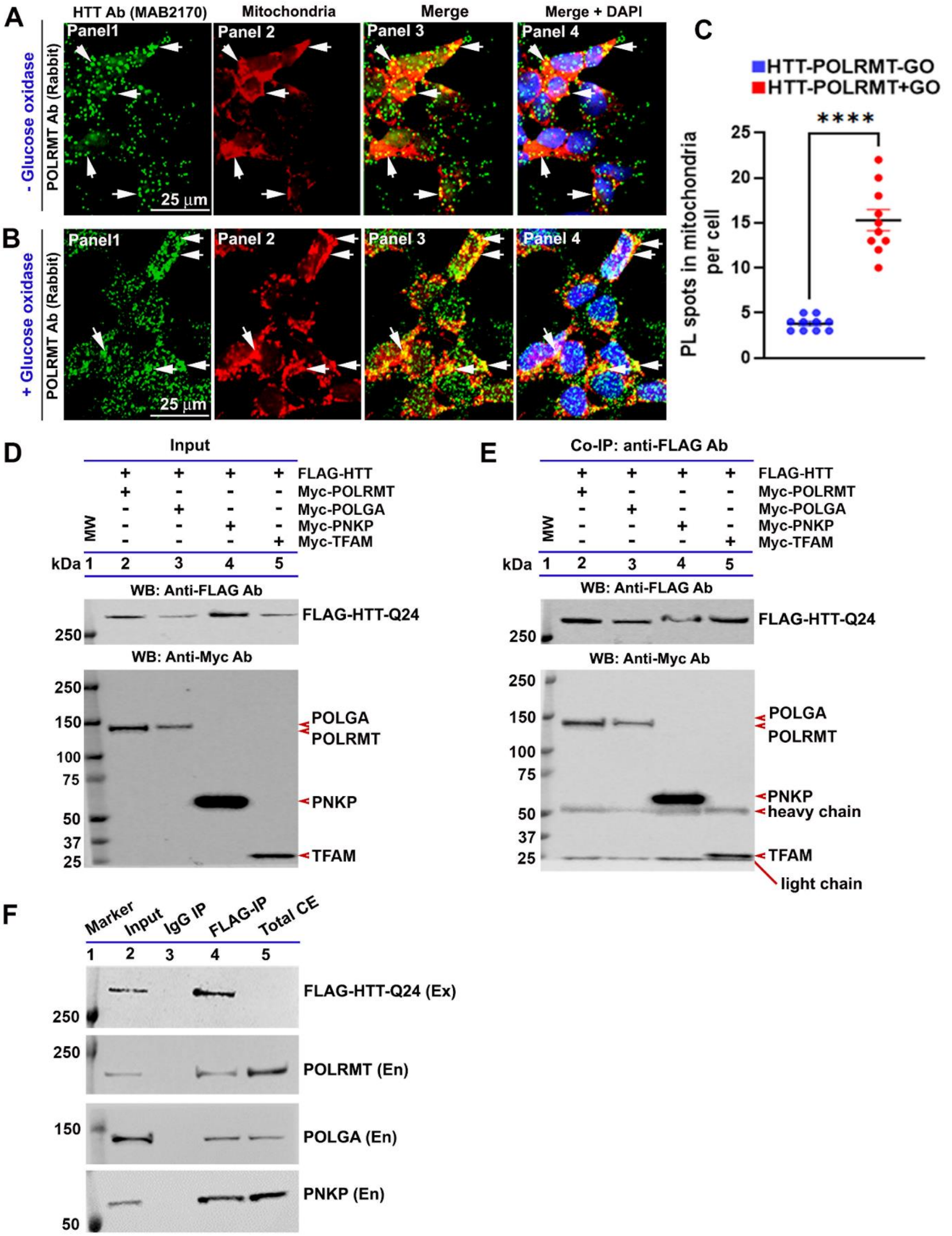

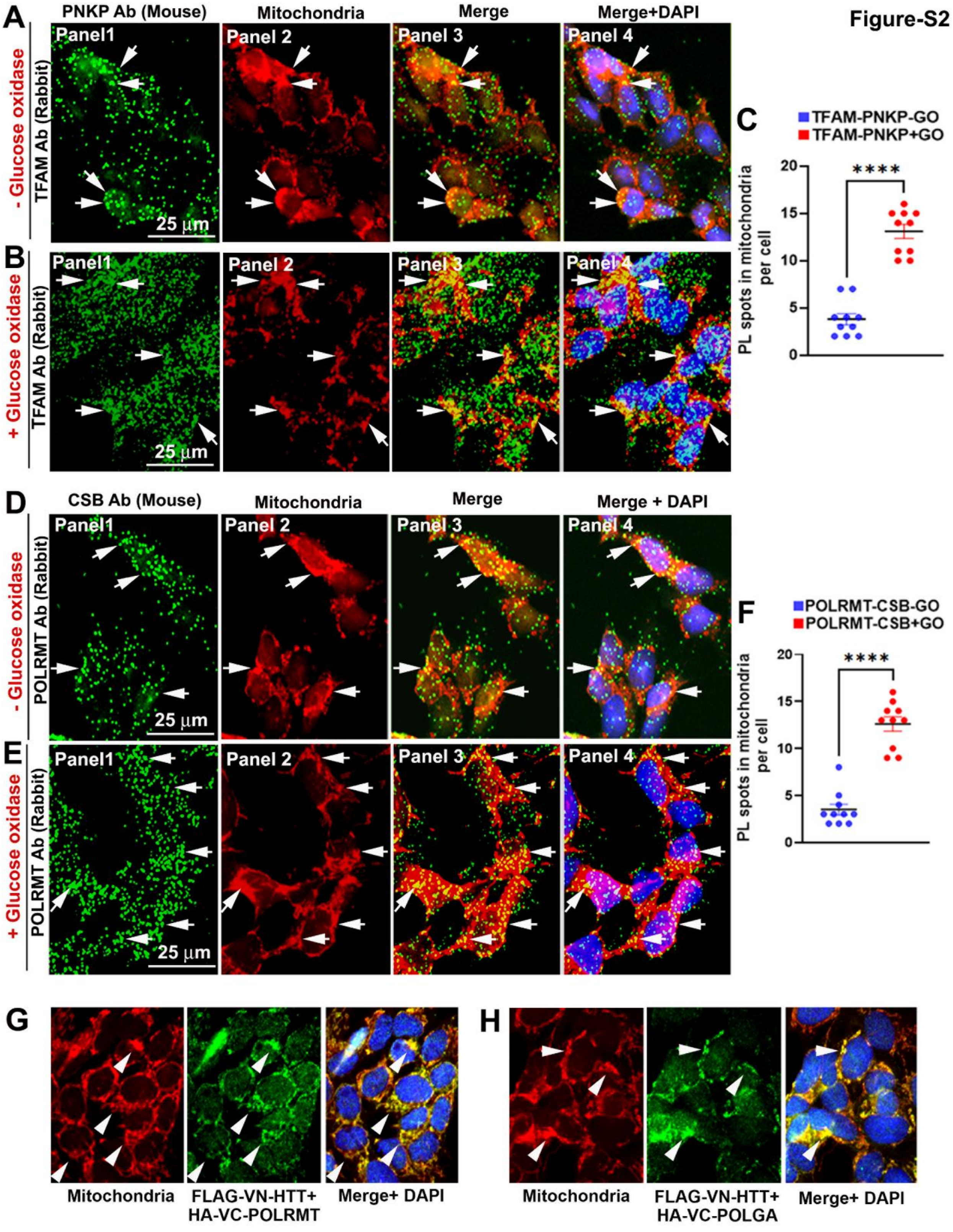

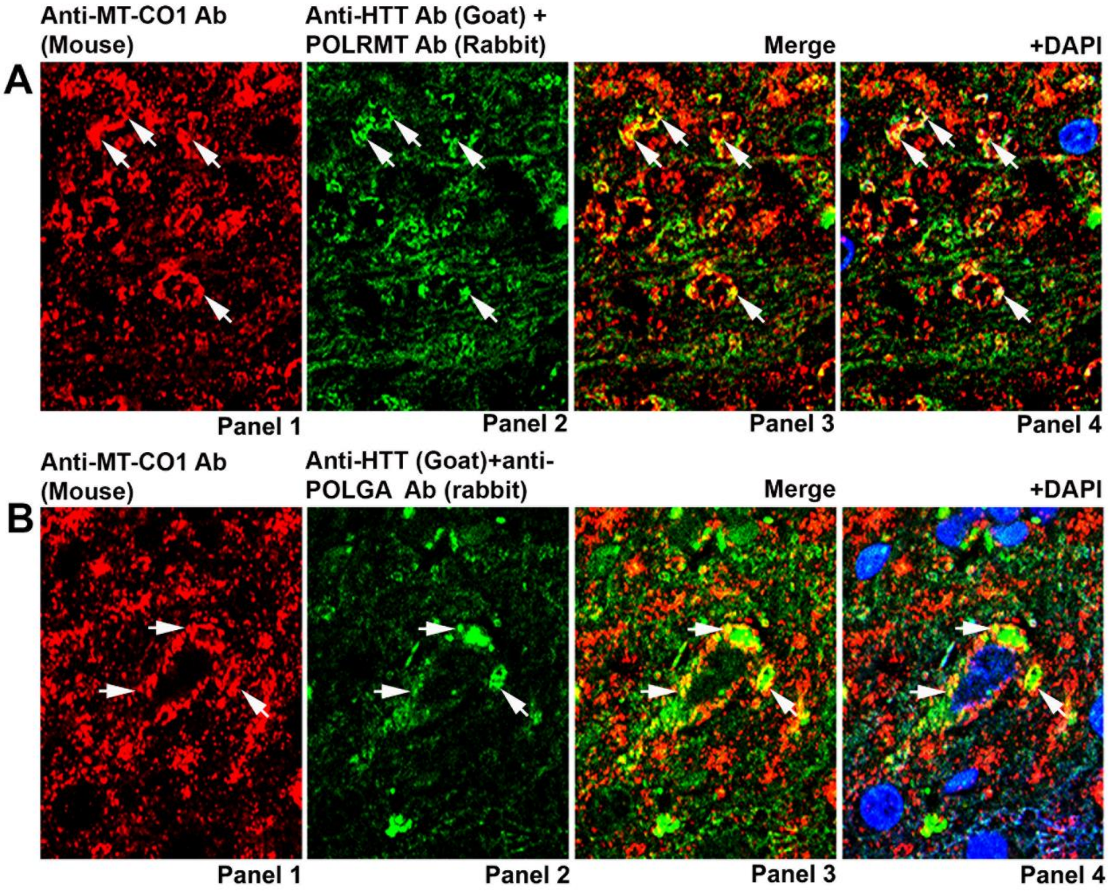

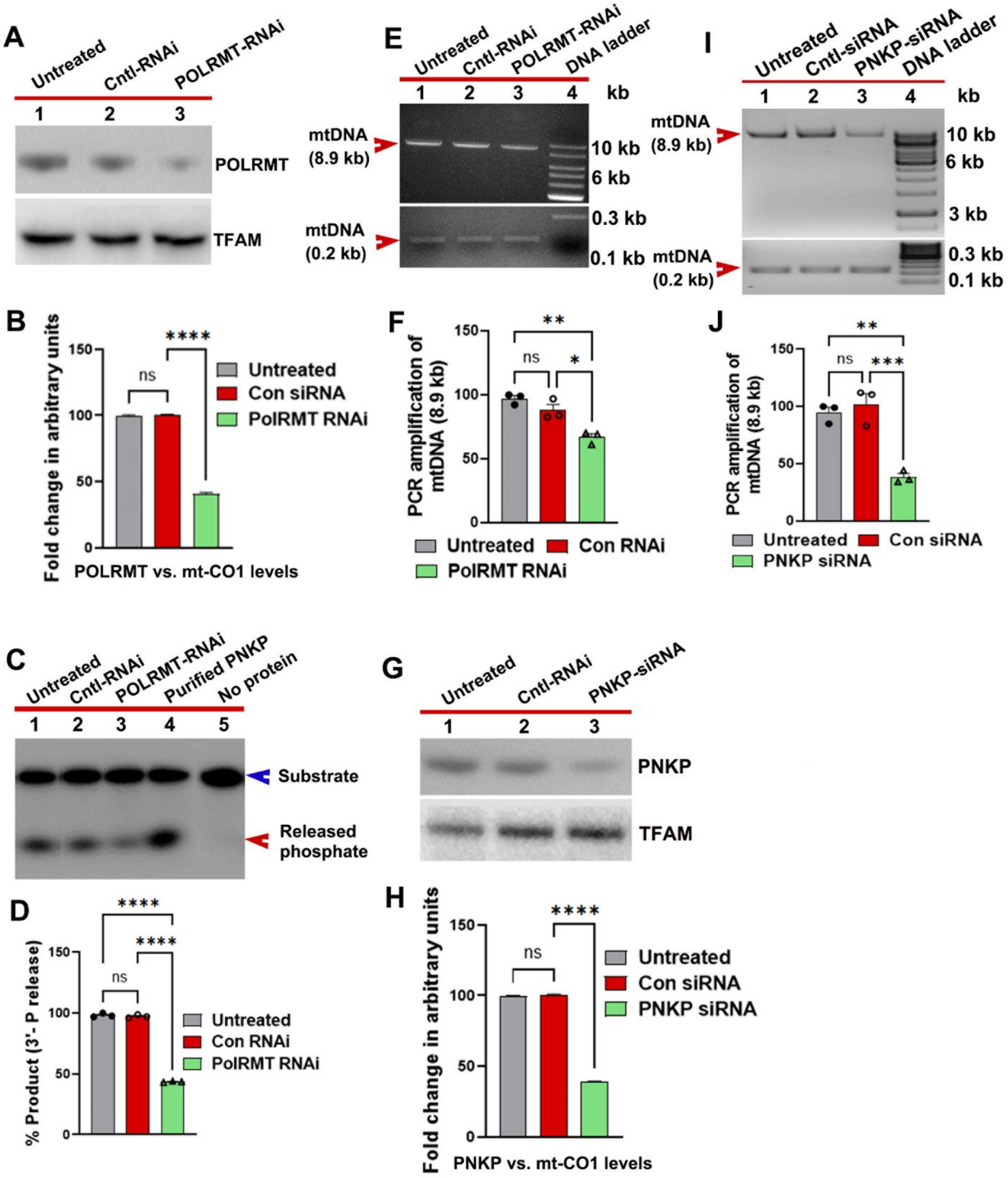

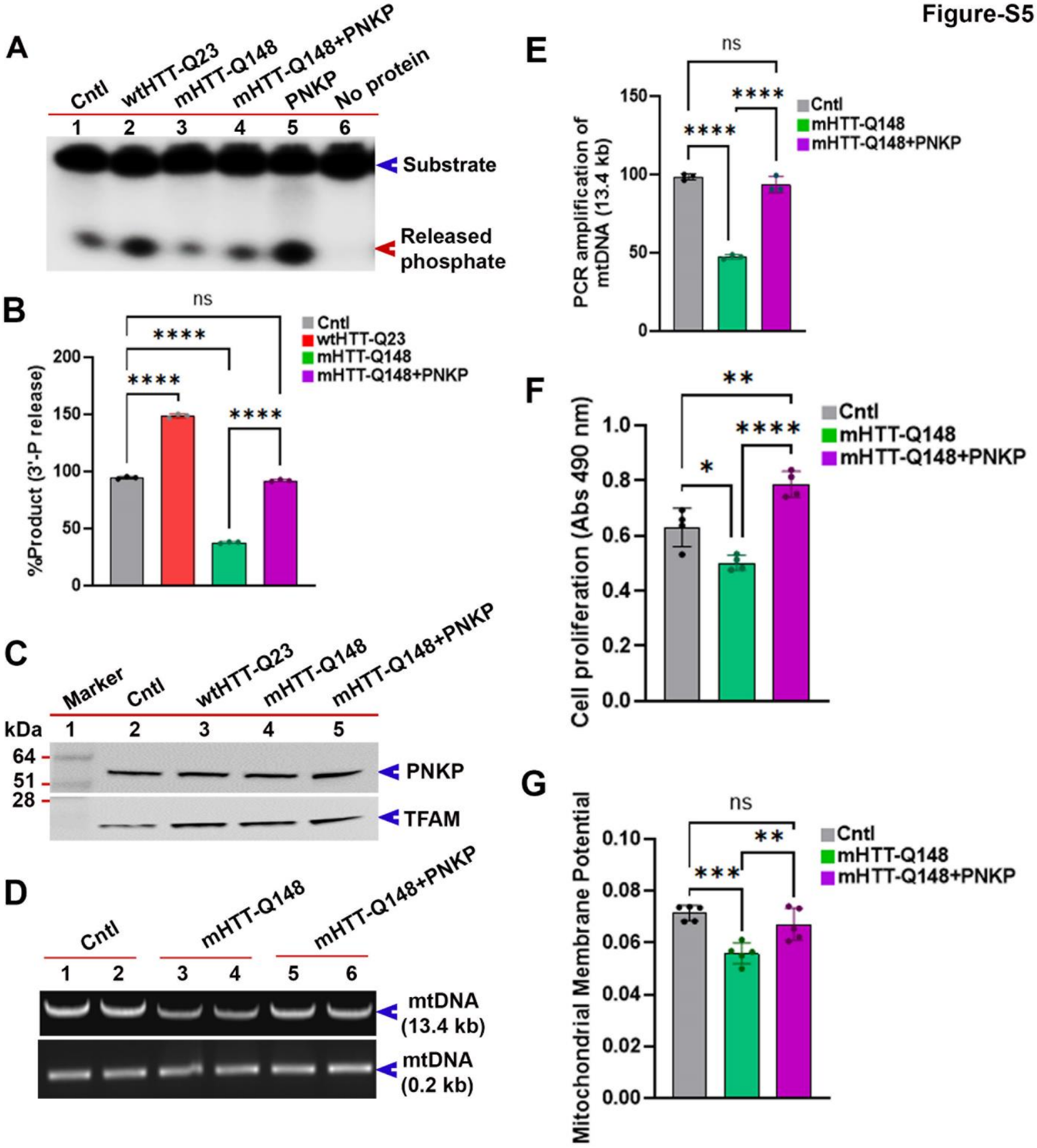

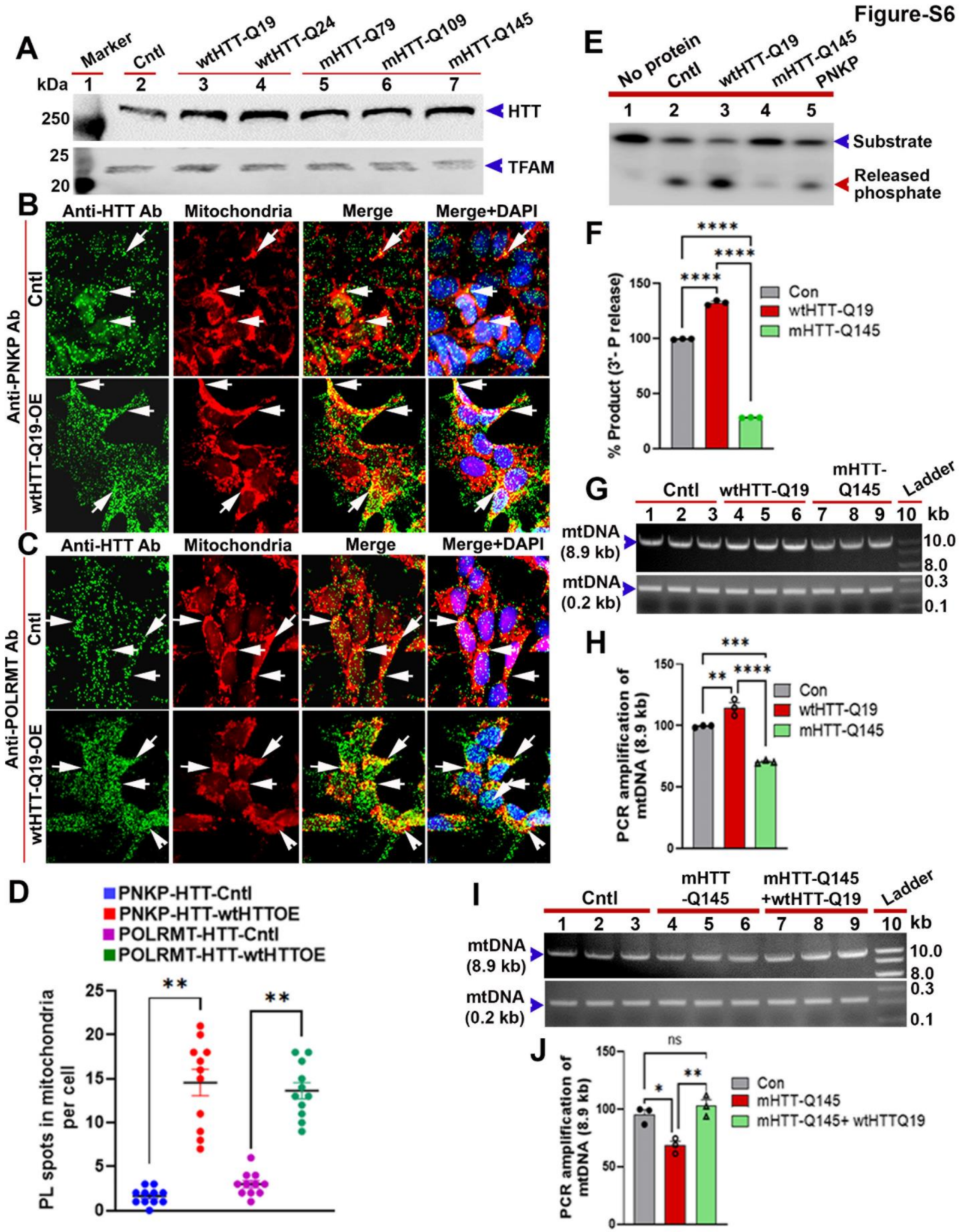

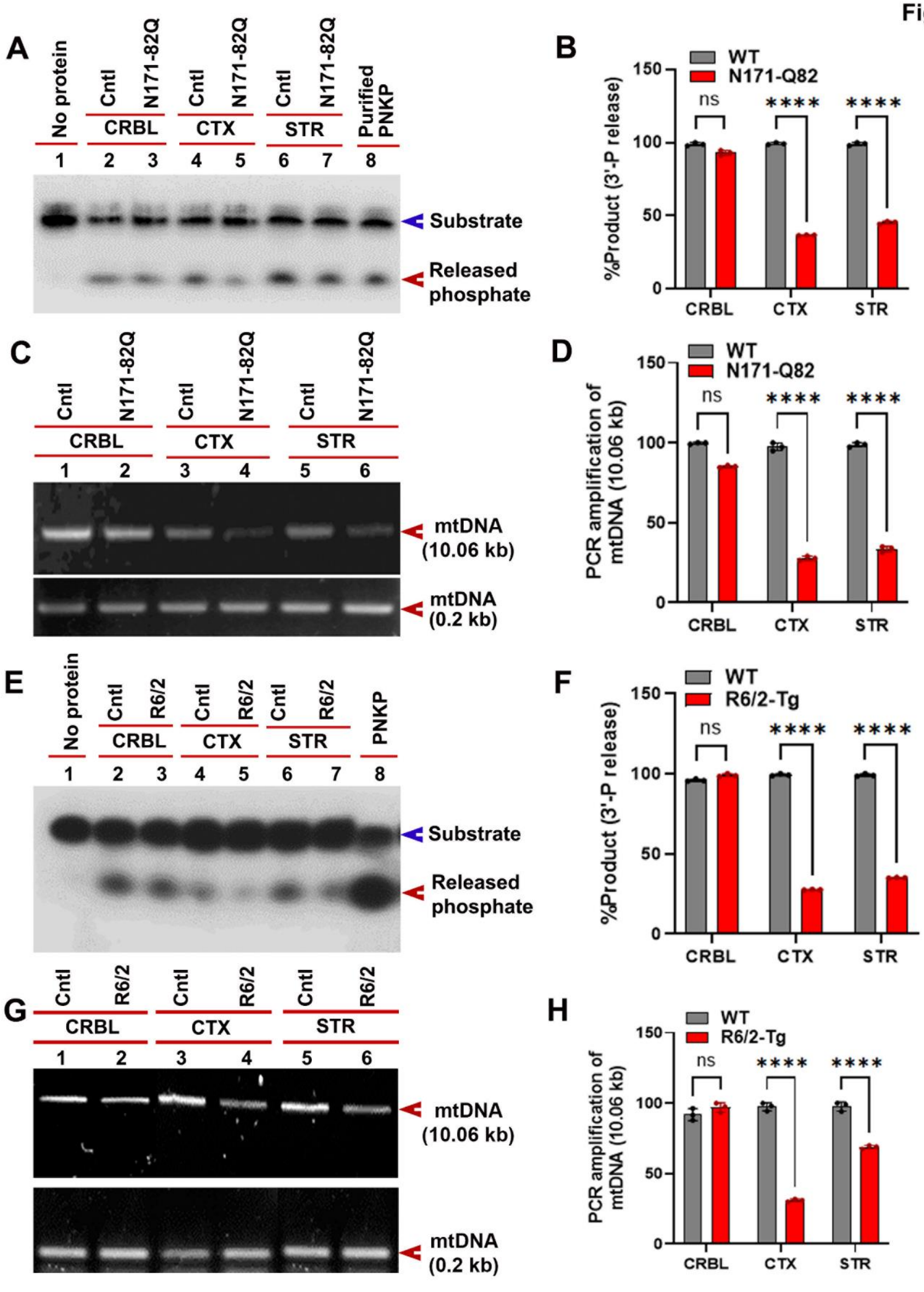
